# Impact of Data Error on Phylogenetic Network Inference from Gene Trees Under the Multispecies Network Coalescent

**DOI:** 10.1101/2025.05.18.654708

**Authors:** Mehrdad Tamiji, Nicolae Sapoval, Luay Nakhleh

## Abstract

Phylogenetic network inference has become an essential tool in evolutionary biology, offering a framework to model complex evolutionary events such as hybridization and horizontal gene transfer. However, a critical but often overlooked challenge is the presence of error in empirical datasets, including sequencing errors, misalignments, and inaccuracies in gene tree estimation. This issue is particularly pressing in the context of phylogenetic networks, which can contain an arbitrary number of parameters and are thus highly susceptible to overfitting. Errors in the input data can lead to artificially inflated network complexity, misrepresenting evolutionary history with non-biological reticulations.

In this study,we systematically examine how different sources of data error influence network inference and show that many widely used methods are vulnerable to these distortions. We find that inaccuracies in gene tree estimation and sequence alignment degrade the reliability of inferred networks. These issues are exacerbated when the number of reticulations that an algorithm can infer exceeds the true number of reticulations in the phylogenetic network. Our analysis underscores the importance of accounting for data error when applying network inference methods and provides practical recommendations for minimizing its impact. By highlighting the vulnerabilities of different approaches and demonstrating how errors propagate through the inference process, we offer practical recommendations for optimizing data processing pipelines. Our findings emphasize the necessity of integrating realistic error models into species network inference methods to enhance their reliability and applicability to real-world biological datasets.

## 1 Introduction

Evolutionary histories are traditionally represented by phylogenetic trees, which model the divergence of species or genes through branching events. In a species tree, branches correspond to speciation events, while in a gene tree, they represent the ancestry of loci descended from a common ancestor. However, several evolutionary processes—including admixture, gene flow, hybridization, hybrid speciation, introgression, and horizontal gene transfer—can give rise to non-tree-like histories. As a result, species trees alone may provide incomplete or inaccurate representations of evolutionary history [1–5]. Forms of gene flow such as introgression and hybrid speciation have been shown to significantly shape evolutionary patterns. When such events occur, a phylogenetic network—a rooted, directed acyclic graph—provides a more accurate model of evolutionary relationships [6]. In this graph, the root represents the most recent common ancestor of all taxa, and internal nodes denote either divergence or reticulation events. Fig. 1 illustrates a phylogenetic network over five taxa, where the blue-colored node inherits genetic material from two distinct ancestral lineages. It is important to note that this layout is purely a visual simplification; graph-theoretically, the underlying structure remains a directed acyclic graph, as used in formal phylogenetic network analysis.

**Figure 1:**
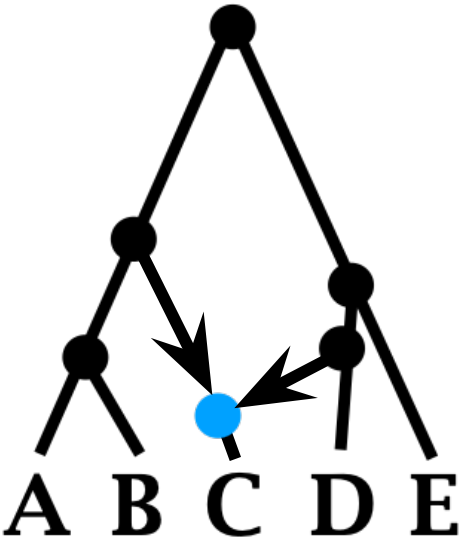
Phylogenetic network of taxa A, B, C, D, and E. The blue node denotes a reticulation event, while the rest of the structure is interpreted in the same way as a phylogenetic tree. While directions are explicitly shown only for the two reticulation edges, all edges are directed away from the root of the network toward the leaves.

The identification of introgression in the presence of incomplete lineage sorting (ILS) motivated the generalization of the multispecies coalescent (MSC) model [7], which defines a probability distribution over gene trees given a species tree and demographic parameters, to the multispecies network coalescent (MSNC) model [8, 9]. This generalization allowed for the development of a wide array of methods for inferring phylogenetic networks under the multispecies coalescent model while also accounting for phenomena such as ILS (e.g., [3, 9–16]). The MSNC allows for modeling evolutionary histories in the presence of both ILS and reticulation, and it enables the derivation of the probability distribution of gene trees on a species network.^1^ Thus, contrasting gene tree distributions under the MSC and MSNC can provide a signal for detecting reticulation and also tease apart the separate roles that ILS and reticulation play.

Methods for inferring species networks under the multispecies network coalescent (MSNC) can be broadly categorized into three classes [17]: summary methods, site-based methods, and coestimation methods. Summary methods estimate networks using inferred gene trees; site-based methods rely on bi-allelic markers; and coestimation methods jointly infer networks and gene trees directly from sequence alignments [18]. Coestimation approaches, such as those described in [12, 14], extend Bayesian frameworks like StarBEAST [19] to phylogenetic networks by sampling from the posterior distribution over species networks and gene trees. However, these methods are computationally intensive and often face convergence and mixing challenges due to the transdimensional nature of network space. Site-based methods, which began with the analytic species tree approach of Bryant et al.[20], were later extended to networks by Zhu et al.[13], and subsequently improved through pseudolikelihood formulations [15] and more efficient inference algorithms [21]. Additional tests for detecting hybridization using bi-allelic markers include the D-statistic [22], its various extensions [23, 24], and invariant-based approaches such as HyDe [25, 26]. Summary methods for the MSNC, originally proposed by Yu et al.[9], employ maximum likelihood estimation from gene tree topologies, with or without branch lengths, and were later extended to a Bayesian framework[11]. To improve scalability, pseudolikelihood approaches were introduced [3, 27], enabling network inference from larger gene tree datasets without the computational burden of full likelihood estimation.

The space of possible network topologies is vast, and their complexity can grow rapidly with the number of taxa and reticulation events. While this complexity can reflect genuine biological processes, it can also arise spuriously due to methodological limitations, gene tree estimation error (GTEE), or model violations. For example, inaccurate or biased gene trees can mislead inference methods into inferring reticulations that do not correspond to actual evolutionary history [28]. Additionally, methods that assume a single gene tree per locus or fail to account for variation in substitution rates across loci may introduce artifacts into the inferred networks [4]. Model misspecification—such as neglecting incomplete lineage sorting (ILS) or overlooking rate heterogeneity—can lead to overfitting, resulting in unnecessary reticulations or incorrect inference of introgression directionality [29]. Notably, recent studies have shown that even fully Bayesian methods, which are theoretically robust, may be misled under rate heterogeneity unless substitution rate variation is explicitly modeled [28, 30]. These issues underscore the importance of statistical rigor in model formulation and biological interpretation of results. Without such diligence, species network inference methods risk producing overly complex networks that reflect methodological artifacts rather than genuine evolutionary signals. In particular, errors introduced during sequencing and multiple sequence alignment can exacerbate these challenges, as they may distort site patterns, mislead gene tree inference, and ultimately inflate the apparent complexity of the inferred networks [31].

In this study, we examined how data errors—specifically sequencing errors and indels—affect species network inference from gene tree estimates under the multispecies network coalescent (MSNC) model. We found that indels substantially increase alignment error and gene tree estimation error (GTEE), particularly in shorter sequences, leading to noticeable shifts in gene tree distributions. Despite these challenges, both maximum likelihood and maximum pseudo-likelihood methods performed well when the number of allowed reticulations matched the true value. However, allowing more reticulations led to overfitting, especially under error, while Bayesian inference via MCMC showed greater robustness due to prior regularization. Although none of the methods could automatically recover the true number of reticulations, tracking changes in (pseudo-)likelihood helped identify a reasonable model complexity, with pseudo-likelihood showing sharper declines in gain and a lower tendency to overfit. These results suggest that current methods—particularly pseudo-likelihood inference—are reasonably robust to moderate data error and can support informed model selection while remaining computationally efficient.

The rest of the chapter is organized as follows. Section 2 provides a brief overview of phylogenomic network inference, a summary of the three network inference methods we study, a detailed description of the simulation setup used to assess the impact of error, and finally information on data and code availability. In Section 3, we present our empirical findings, highlighting how different types of errors affect inference accuracy across methods. Section 4 offers a discussion of the implications of our results, emphasizing practical recommendations for data analysis and suggesting directions for improving model robustness in the presence of error. Afterwards, we propose a set of guidelines for biologists to aid in the design and interpretation of phylogenetic analyses and outline directions for future research to address open challenges and expand upon our findings.

## 2 Materials and Methods

This section outlines the methods used to study the impact of data error on species network inference, including a review of inference models, the simulation setup, and details on data and code availability.

### 2.1 Species network inference under the multispecies network coalescent: A brief overview

A typical phylogenomic inference pipeline for species networks is illustrated in Fig. 2. To infer a species network on a set 𝒳 of species, sequence data are obtained from multiple loci. For the phylogenomic methods studied in this chapter, we assume the loci are independent and recombination-free. The sequences for each locus are then aligned, and from a modeling and inference perspective, the data are represented as a set *S* = {*S*_1_, …, *S*_*m*_} of multiple sequence alignments (MSAs) from *m* loci, where *S*_*i*_ denotes the MSA for locus *i*.

**Figure 2:**
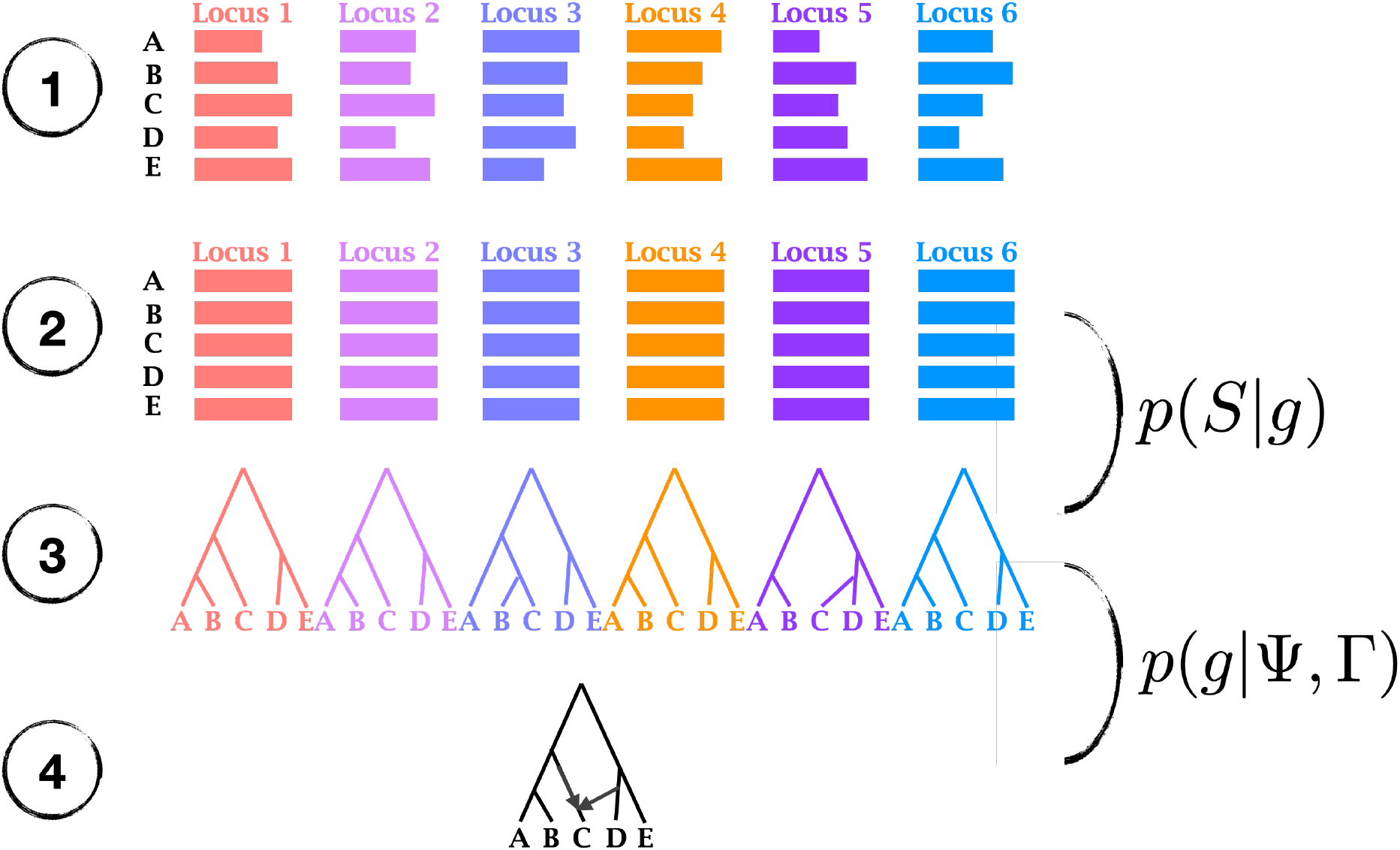
Phylogenomic pipeline of species network inference. (1) Orthologs from five species—A, B, C, D, and E—are obtained across multiple loci (six in this case). (2) For each locus, the sequences are aligned to produce one multiple sequence alignment (MSA) per locus. (3) For many species phylogeny inference methods, the input consists of gene trees, typically one per locus. For these methods, a gene tree is constructed from the MSA of each locus. The methods examined in this chapter assume that the gene trees are also rooted. (4) A species network is inferred from the set of gene trees. In likelihood-based approaches, the MSA *S* and gene tree *g* of each locus are related by the likelihood function *p* (*S* | *g*), and each gene tree *g* is related to the species network Ψ and its inheritance probabilities by the likelihood function *p* (*g* | Ψ, Γ).

Phylogenetic networks have been defined in the literature to refer to various models [32]. In this work, we focus on species networks that are explicit phylogenetic networks, i.e., ones that explicitly model the evolutionary history of a set of taxa; Fig. 2. In this context, a *species network* Ψ on a set 𝒳 of taxa is a rooted, directed, acyclic graph with a set *V* of nodes (or vertices) and a set *E* of edges (or branches). Except for the root node, which has no parents, each node in *V* has either one or two parents. Nodes with two parents are called *reticulation nodes*, and all other nodes—including the root—are referred to as *tree nodes*. The edges connecting a reticulation node to its two parents are termed *reticulation edges*, while all other edges are called *tree edges*. The leaf nodes (nodes with no children) are uniquely labeled by the taxa in 𝒳. The species network in Fig. 2 shows the evolutionary history of the set 𝒳 = {*A, B, C, D, E*} and contains one reticulation node, which is the endpoint of two arrows denoting the reticulation edges. Notably, if one of these arrows is removed, the resulting phylogeny takes the shape of a tree, illustrating that a phylogenetic tree is a special case of a species network—one that contains no reticulation nodes.

When a species network Ψ is augmented with branch lengths in coalescent units and inheritance probabilities (every two reticulation edges are assigned nonnegative real numbers that sum up to 1), the model specifies the multispecies network coalescent, which defines a probability distribution over gene trees [9]. The mass and density functions of gene trees given a species network, its branch lengths, and inheritance probabilities were derived in [8, 9, 33].

Given a model consisting of a species network Ψ (including its topology and branch lengths) and inheritance probabilities Γ, the posterior distribution over the model parameters is defined as

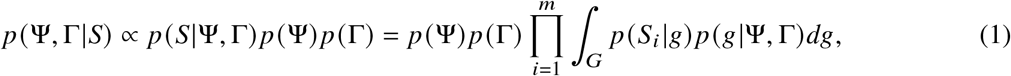

where *p* (*S*_*i*_ |*g*) denotes the probability of the sequence alignment *S*_*i*_ given a particular gene tree *g* [34], and *p* (*g* |Ψ, Γ) represents the density of the gene tree (including its topology and branch lengths) given the model parameters [9].

If the set *G* of gene trees has been estimated for the *m* loci, with gene tree *g*_*i*_ estimated from sequence alignment *S*_*i*_, we can treat *G* as the data. The posterior distribution of the model is then given by

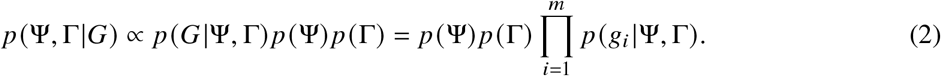

Furthermore, under this formulation, the maximum likelihood estimate (MLE) of the species network and its parameters is given by

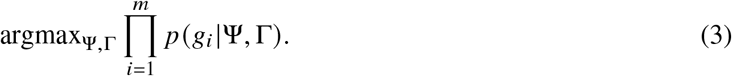

Calculating the likelihood function 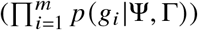 is computationally very expensive [35], which has motivated the development of a maximum pseudo-likelihood approach [27] and a pseudo-likelihood-based Bayesian inference method [36].

### 2.2 Inference methods

In this work, we focus on three methods for phylogenomic network inference from gene trees that are implemented in `PhyloNet` [37, 38]: maximum likelihood inference (ML) under the MSNC [9], maximum pseudo-likelihood inference (MPL) under the MSNC [27], and pseudo-likelihood-based Bayesian MCMC inference (PL MCMC) under the MSNC [11, 36]. All of these methods take as input a set of rooted gene trees. Details of these methods and their parameters are provided in the paragraphs below.

#### InferNetwork ML

is a likelihood-based method designed to infer species networks that model reticulate evolutionary histories while accounting for incomplete lineage sorting (ILS) [9]. The method operates by maximizing the likelihood (Eq. 3) of observed gene trees under the MSNC. The process involves a heuristic search over the space of species networks with an upper bound on the maximum number of allowed reticulations. By default, the search begins with the species tree that minimizes the number of deep coalescences [39].

The script in Listing 1 provides a sample command for running the `InferNetwork ML` method with *M* gene trees, a maximum of three reticulations, and utilizing a single CPU core (via the `-pl 1` flag). The output of the command provides an estimated species network and its likelihood.

**Listing 1:**
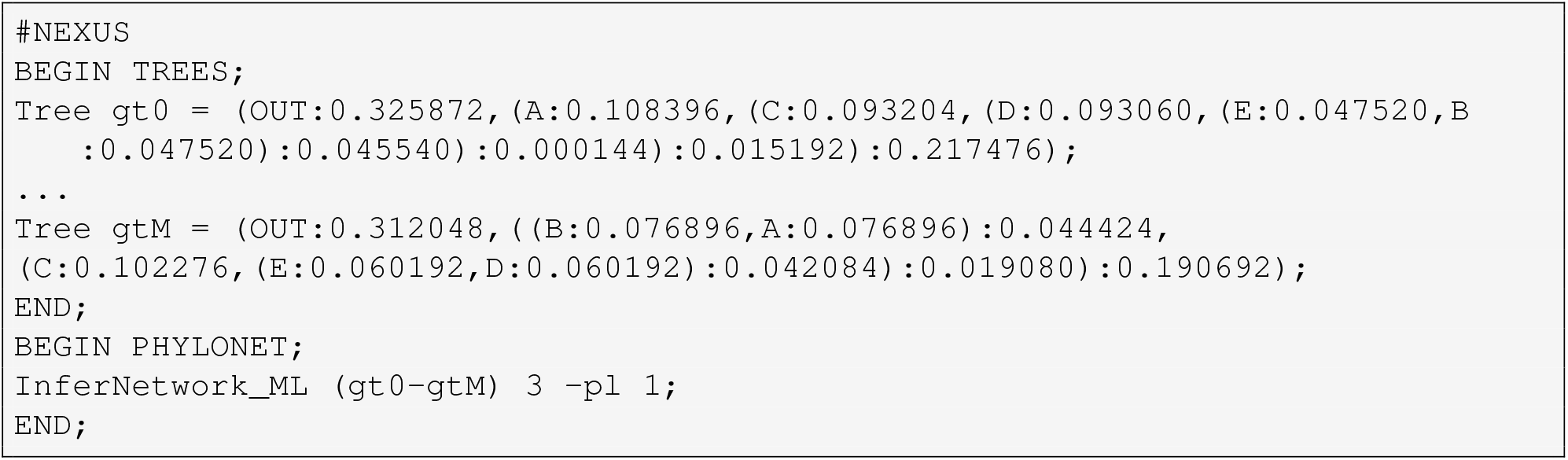
Example NEXUS file for inferring species networks using `InferNetwork ML`.

#### InferNetwork MPL

is a pseudo-likelihood-based method designed for inferring phylogenetic networks while accounting for incomplete lineage sorting (ILS) [27]. Unlike `InferNetwork ML`, which directly maximizes the full likelihood of the network, `InferNetwork MPL` approximates the likelihood using a pseudo-likelihood function based on the distribution of the rooted triples of taxa.

The script in Listing 2 provides an example command for running the method in `PhyloNet`. Similar to `InferNetwork ML`, this command specifies *M* gene trees, sets a maximum of three reticulations, and runs the computation using a single CPU core (via the `-pl 1` flag). The output of the command includes an estimated species network and its associated pseudo-likelihood.

**Listing 2:**
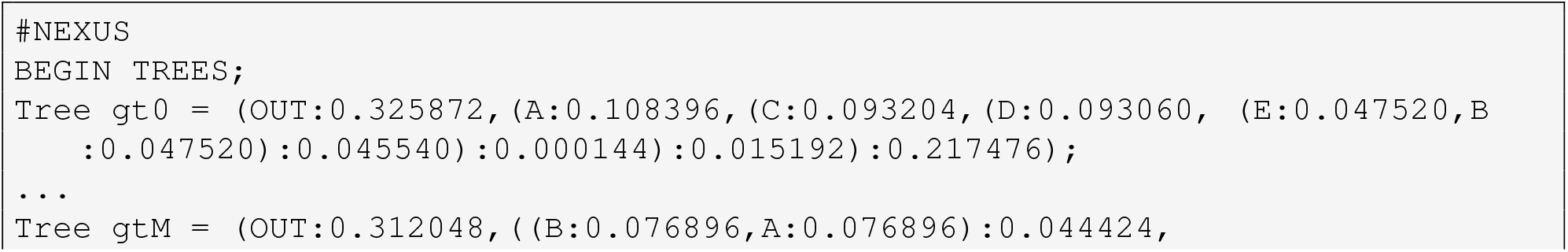
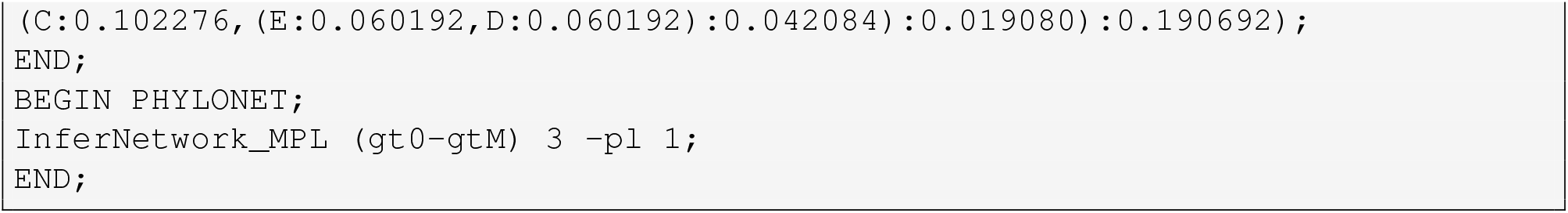
Example NEXUS file for inferring species networks using `InferNetwork MPL`.

#### MCMC GT Pseudo

`MCMC GT Pseudo` is a Bayesian inference method for species networks that leverages pseudo-likelihood [11, 36]. Unlike `InferNetwork ML` and `InferNetwork MPL`, which maximize the likelihood and pseudo-likelihood, respectively, `MCMC GT Pseudo` employs a Markov chain Monte Carlo (MCMC) framework to sample from the posterior distribution of species networks.

The script in Listing 3 provides an example command for running the method in `PhyloNet`. This command specifies *M* gene trees, sets a maximum of three reticulations, and utilizes a single CPU core. The `-pseudo` flag indicates the use of pseudo-likelihood during sampling. Additional parameters for MCMC sampling are also specified: `-cl` sets the total chain length, `-bl` defines the number of burn-in iterations (i.e., discarded samples to ensure convergence), and `-sf` determines the sampling frequency. The output of the command provides all sampled species networks, along with summary statistics for all topologies in the 95% credible set, including the MAP and average branch length estimates.

**Listing 3:**
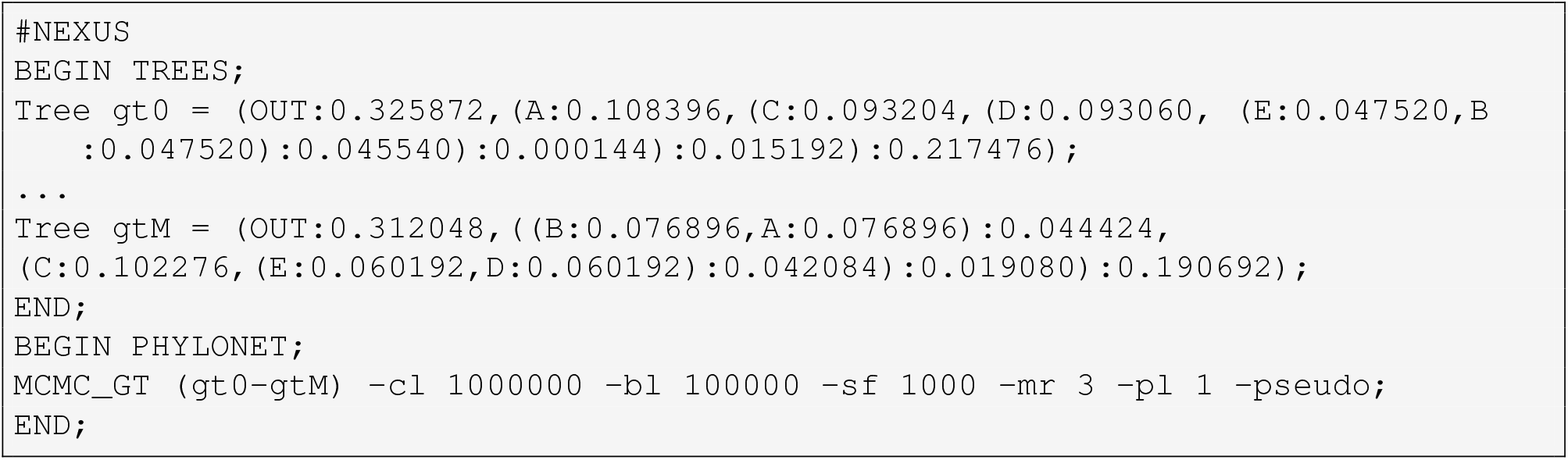
Example NEXUS file for inferring species networks using `MCMC GT Pseudo`.

### 2.3 Simulation setup

To comprehensively evaluate the impact of various modes of error on species network inference, we designed an extensive experimental setup (Fig. 3) that enables the assessment of individual contributions from each potential error source in a progressively more realistic manner under a variety of parameter settings.

**Figure 3:**
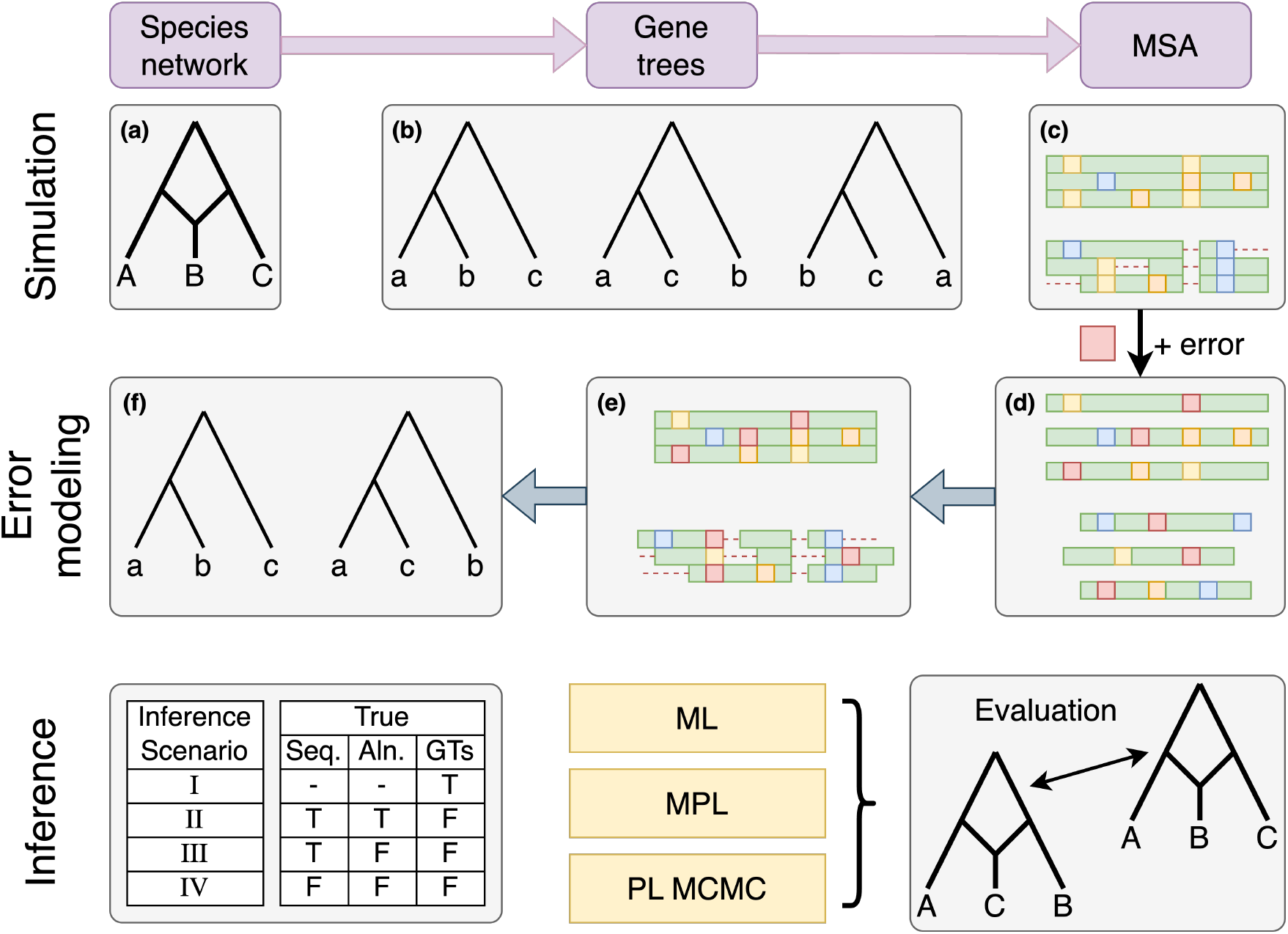
Schematic representation of the experimental setup. **(a)** Model species networks are designed by hand. **(b)** Gene trees are simulated on the species networks under the MSNC model. **(c)** Alignments with and without indels are simulated for each gene tree using INDELible. **(d)** Sequence error is added to individual sequences in the alignment. **(e)** Sequences are re-aligned using MAFFT to obtain empirical MSAs. **(f)** Gene trees are inferred under ML with IQ-TREE. In total, four scenarios (I–IV) are analyzed: (I) inference from true gene trees, (II) inference from gene trees inferred from true alignments, (III) inference from gene trees inferred from inferred alignments of true sequences, and (IV) inference from gene trees inferred from alignments of sequences with error. For species network inference, we ran three methods: maximum likelihood (ML), maximum pseudo-likelihood (MPL), and Bayesian MCMC with pseudo-likelihood (PL MCMC).

We used three model networks in our simulation study, each with a different number of reticulations (Fig. 4).

**Figure 4:**
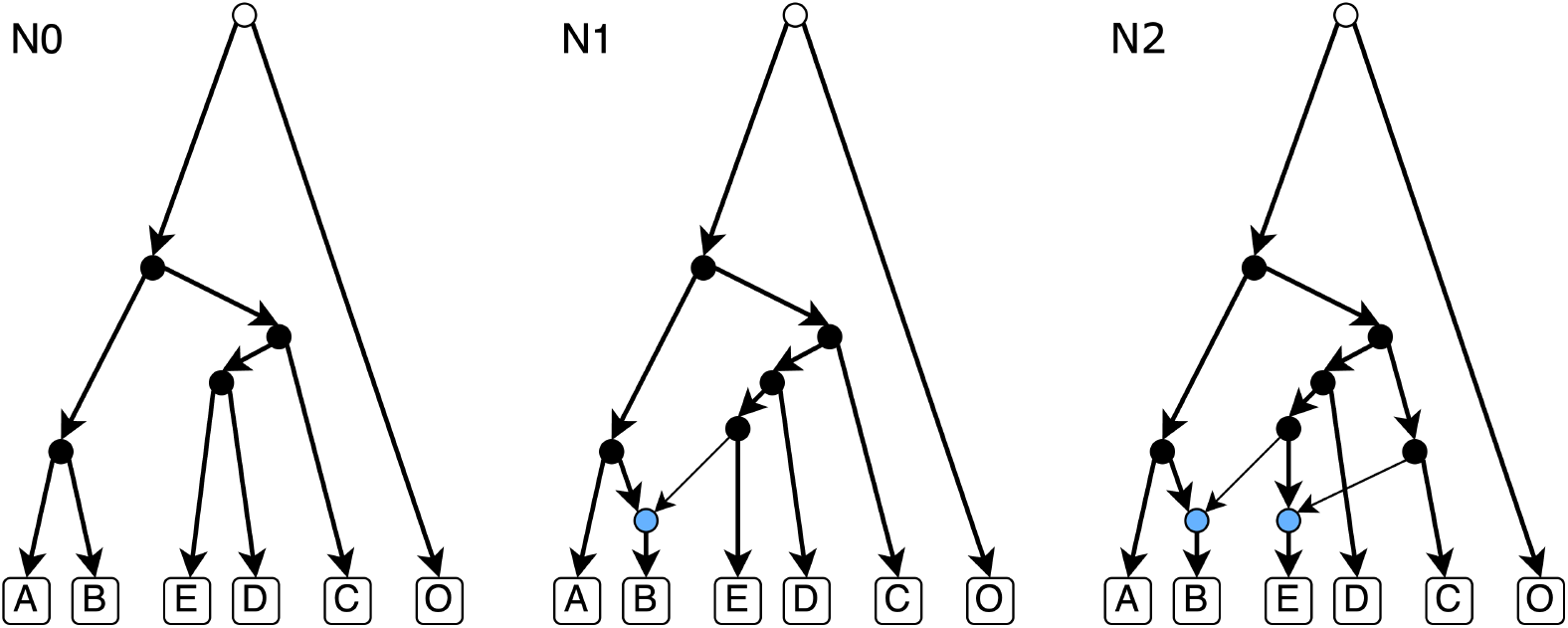
Schematic representation of three species networks with varying numbers of reticulations. The three networks shown—N0, N1, and N2—on five ingroup taxa (A, B, C, D, and E), one outgroup (O), and 0, 1, and 2 reticulation nodes, respectively. All networks have a total height of 16 coalescent units, with the branch from the root to the most recent common ancestor of the ingroup spanning 10 coalescent units. Reticulation nodes are shown in blue, and thinner arrows indicate reticulation edges with an inheritance probability of 0.3.

The corresponding extended Newick representations are listed in Table 1.

**Table 1:**
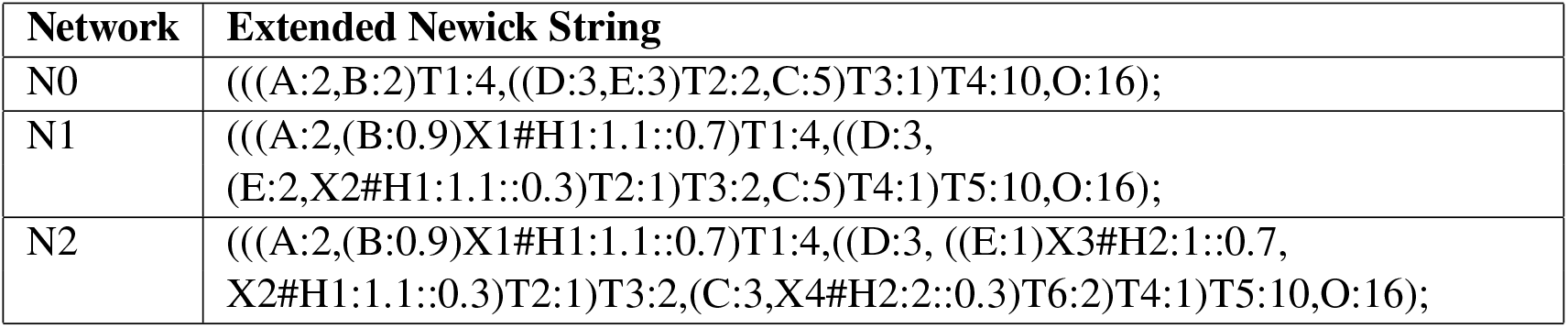
Extended Newick strings for the three model networks used in the simulations.

Namely, for each of the three model networks (labeled N0, N1, and N2 in Fig. 4, where the index indicates the number of reticulations in the network), we simulated 100, 250, and 500 gene trees under the MSNC using the `SimGTInNetwork` wrapper in `PhyloNet` for the `ms` [40] command. The branch lengths of the simulated gene trees generated by `ms` are in coalescent units per generation. We converted these to mutation units by applying a scaling factor of 0.018 when simulating the alignments.

An example command for generating 100 gene trees for a species tree is provided in Listing 4.

**Listing 4:**
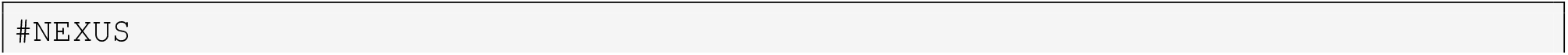
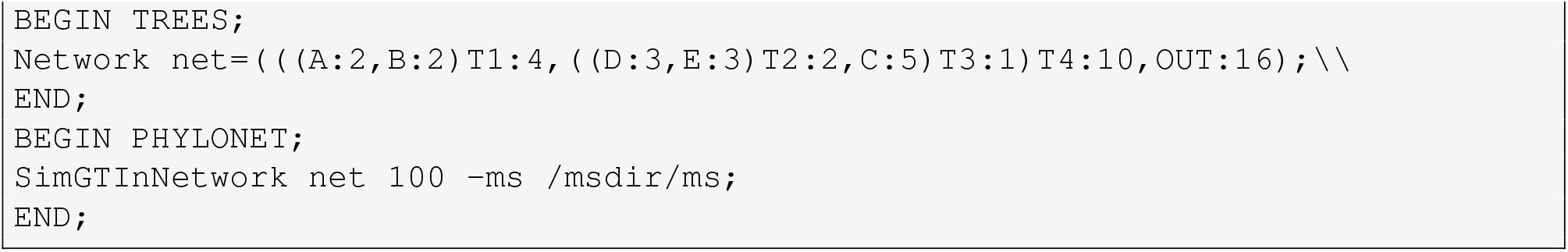
Example NEXUS file for simulating gene trees under the MSNC in `PhyloNet`.

For each simulated gene tree, we simulated an MSA under the GTR model using `INDELible` [41]. We considered three alignment lengths: 250, 500, and 1000. For all alignments, we used the following common GTR parameters: (a) substitution rates AC=0.2173, AG=0.9798, AT=0.2575, CG=0.1038, CT=1.0, and GT=0.207; (b) base frequencies A=0.2112, C=0.2888, G=0.2896, and T=0.2104. For simulating indels, we used a power-law (POW) distribution with *α* = 1.5 and a maximum indel length of 5. We considered three indel rate values: 0, 0.05, and 0.1. Together, these parameters define our ground truth simulation data for gene trees and MSAs (Fig. 3(b,c)).

To use `INDELible`, one needs to create a `control.txt` file and then run `INDELible` using that file. Listing 5 shows an example `control.txt` configuration. The output is a single FASTA alignment of 1000 nucleotides, which may include gaps due to indels.

**Listing 5:**
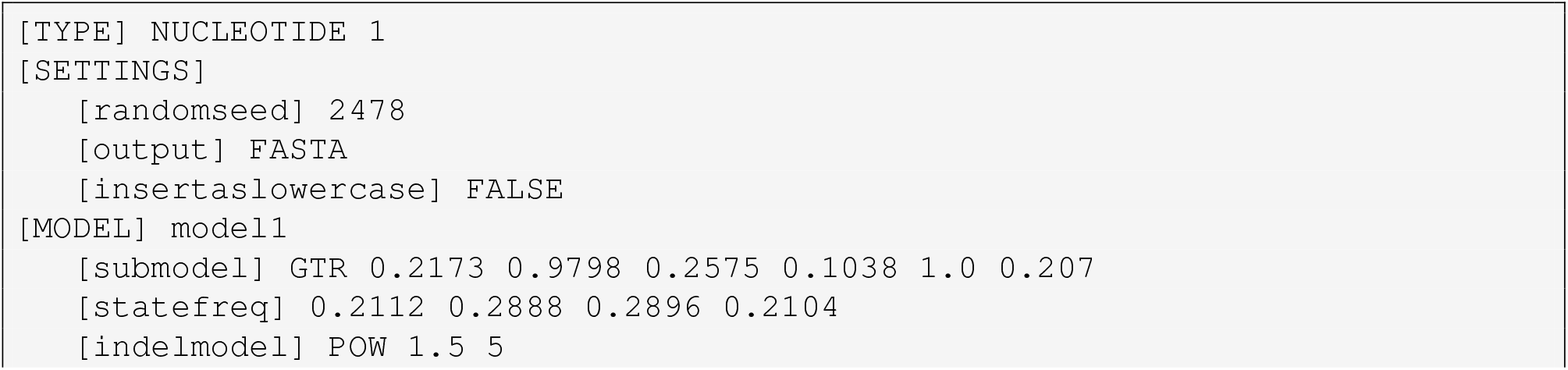
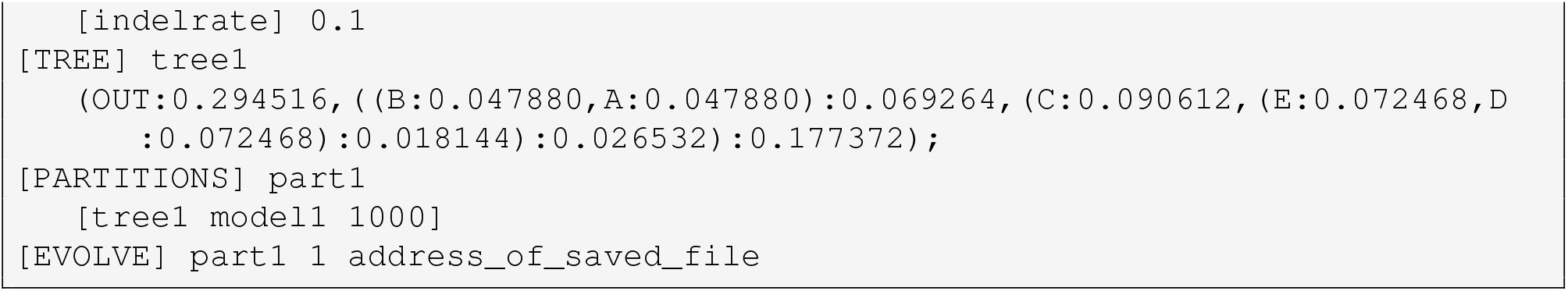
An example `control.txt` file for sequence generation in `INDELible`.

In order to mimic potential sequencing and/or assembly errors, we simulated sequence errors (Fig. 3(d)) in the simulated gene sequences by introducing random nucleotides with probabilities of 0 (no error), 0.01, and 0.1. Every erroneous nucleotide had the uniform probability of being any of the four nucleotides (i.e., once a nucleotide was selected for error introduction, it was replaced with a different nucleotide with probability 0.75).

We considered progressively more complex scenarios (Fig. 3(I-IV)), starting with inference from true gene trees and ending with inference from gene trees reconstructed from alignments of erroneous sequences. For MSA inference, we used MAFFT [42], specifically the iterative refinement algorithm for global multiple sequence alignment (`G-INS-i`, default). An example alignment command is shown in Listing 6.

**Listing 6:**
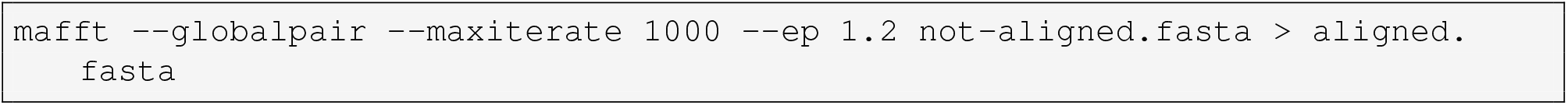
MAFFT command used for sequence alignment.

To infer gene trees from true alignments (scenario II), inferred alignments (scenario III), and inferred alignments over erroneous sequences (scenario IV), we used `IQ-TREE` [43] with the ModelFinder Plus (MFP) option. Since `IQ-TREE` generates unrooted trees, we subsequently rooted the inferred gene trees using the designated outgroup taxon. A sample command for running `IQ-TREE` with bootstrap is shown in Listing 7.

**Listing 7:**
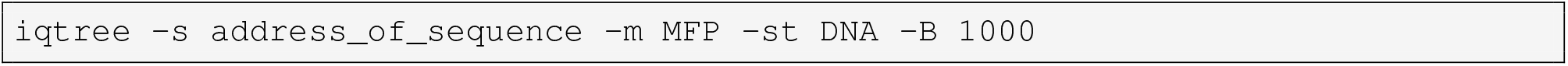
IQ-TREE command used for gene tree inference.

### 2.4 Data and code availability

All simulated data produced for our analyses have been archived and are available on Zenodo [44]. All control files, extended Newick strings for the species networks, and scripts used to generate data, run analyses, and plot results are available on GitHub [45]. `PhyloNet` is open source and available on GitHub [46].

## 3 Results

This section presents our experimental findings, beginning with an analysis of the characteristics of the simulated data. We then examine the likelihood trajectories across different error scenarios. Subsequent subsections explore the distributions of MCMC samples, the topological accuracy of inferred networks, and the computational costs associated with the inference methods.

### 3.1 Characteristics of the simulated data

First, we summarized the characteristics of the inferred MSAs and gene trees in order to assess the levels of error introduced by standard inference procedures across varying indel rates in the true sequences and rates of erroneous nucleotides.

*Multiple sequence alignment quality* was quantified using the sum-of-pairs (SP) score, computed as the average (over pairwise alignments induced by the MSA) ratio between correctly aligned base pairs in the inferred alignment and the total number of aligned base pairs in the reference alignment. The results are shown in Fig. 5.

**Figure 5:**
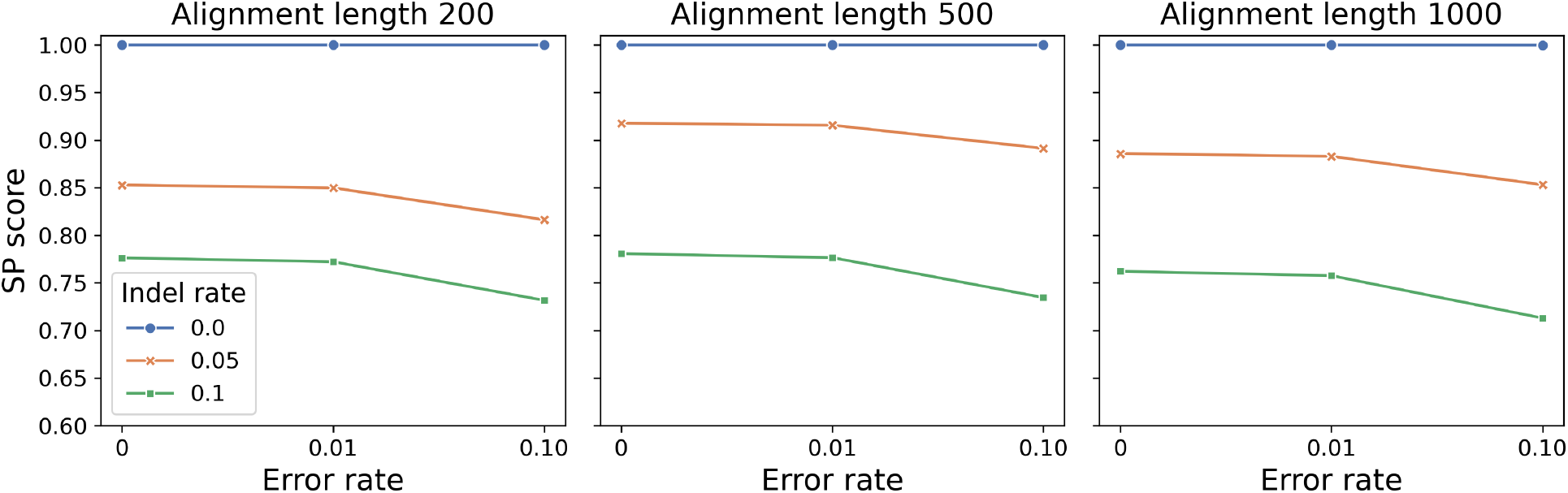
Alignment quality for varying indel and error rates. Variability in the average SP scores (y-axis) of the alignments based on the indel rate (indicated by color), error rate (x-axis), and alignment length (subplots).

We observe that when the indel rate is set to 0, i.e., the simulated alignment has no indels, the average SP score of all alignments remains close to 1 regardless of the error rate. Furthermore, we note that the indel rate alone has a greater impact on alignment quality, resulting in a mean SP score drop of 0.114 from an indel rate of 0 to 0.05, and a drop of 0.227 from 0 to 0.1. In comparison, increasing the error rate from 0 to 0.1 has no impact on indel-free alignments, and results in an average SP score decrease of 0.032 for alignments with an indel rate of 0.05, and 0.047 for those with an indel rate of 0.1 (Fig. 5). Notably, there is little variation in alignment quality across independent replicates, with the mean standard deviation of SP scores between replicates being 0.0004, a maximum of 0.0012, and a minimum of 0.

*Gene tree estimation error (GTEE)* was quantified by measuring the average normalized Robinson–Foulds (nRF) distance [47] between ground-truth simulated gene trees and gene trees inferred from: error-free true alignments (scenario II), error-free MAFFT alignments (scenario III), and MAFFT alignments of sequences with erroneous nucleotides (scenario IV). The results are shown in Fig. 6.

**Figure 6:**
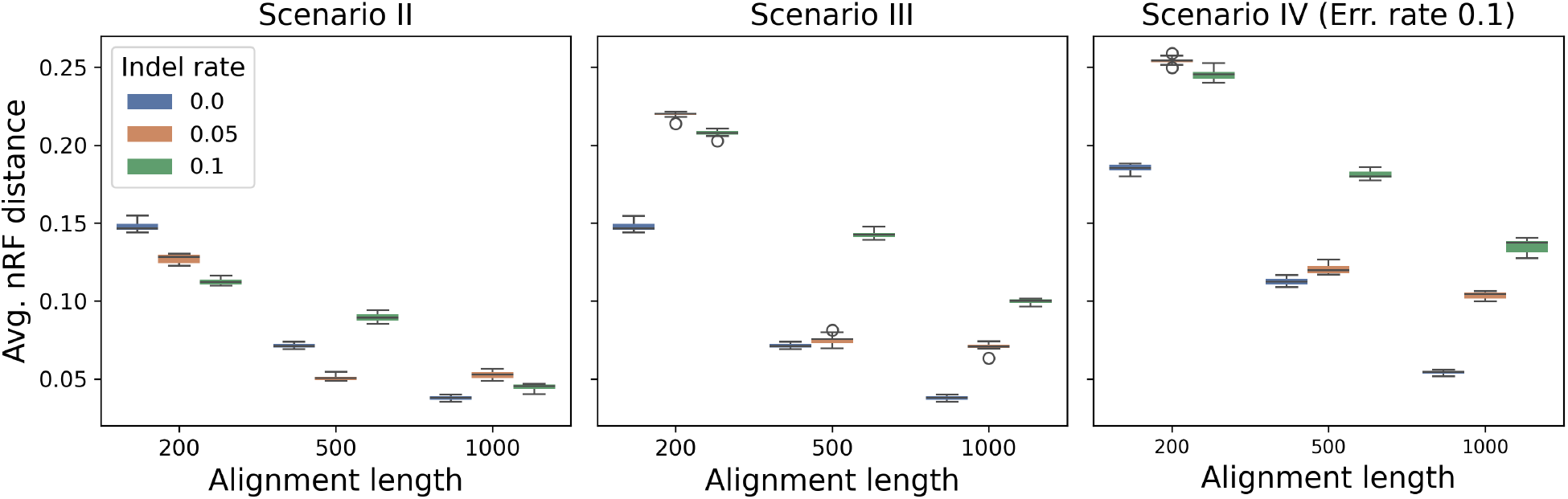
Gene tree estimation error under different data error modes. Boxplots of normalized RF distances (y-axis) between ground-truth gene trees and inferred gene trees for the three scenarios: error-free true alignments (leftmost panel), MAFFT alignments of error-free sequences (middle panel), and MAFFT alignments of sequences with erroneous nucleotides at an error rate of 0.1 (rightmost panel). Alignment length is shown on the x-axis, and colors represent varying indel rates in the ground-truth alignment.

We observe that in all cases, GTEE decreases as alignment length increases. In the case of true alignments (scenario II), indels have a mixed effect on gene tree inference accuracy, while in scenarios III and IV, the presence of indels consistently increases GTEE. Consistent with the alignment accuracy results, indels generally have a greater impact on GTEE than the error rate alone.

We calculated the *Kullback–Leibler (KL) divergence* between the empirical distributions of inferred and true gene tree topologies to assess the impact of data error on the inferred gene tree distribution. This measure is particularly relevant because species network inference under the MSNC utilizes the distribution of gene trees. Thus, distortions in this distribution—caused by alignment error, sequencing error, or indels in our

study—can directly affect the accuracy of network inference. For each replicate, we enumerated the unique topologies across gene trees and computed their empirical frequencies. Let *P* denote the true distribution and *Q* the inferred one. The KL divergence is then computed as:

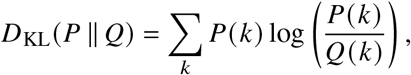

where *k* ranges over all observed topologies. To handle cases where *Q* (*k*) = 0, we applied Laplace smoothing by adding a small constant (*ϵ* = 1*e* − 5) to all gene tree frequencies prior to calculating the divergence.

Fig. 7 shows KL divergence values across increasing alignment lengths (x-axis) for three evolutionary models (rows) under three different scenarios (columns).

**Figure 7:**
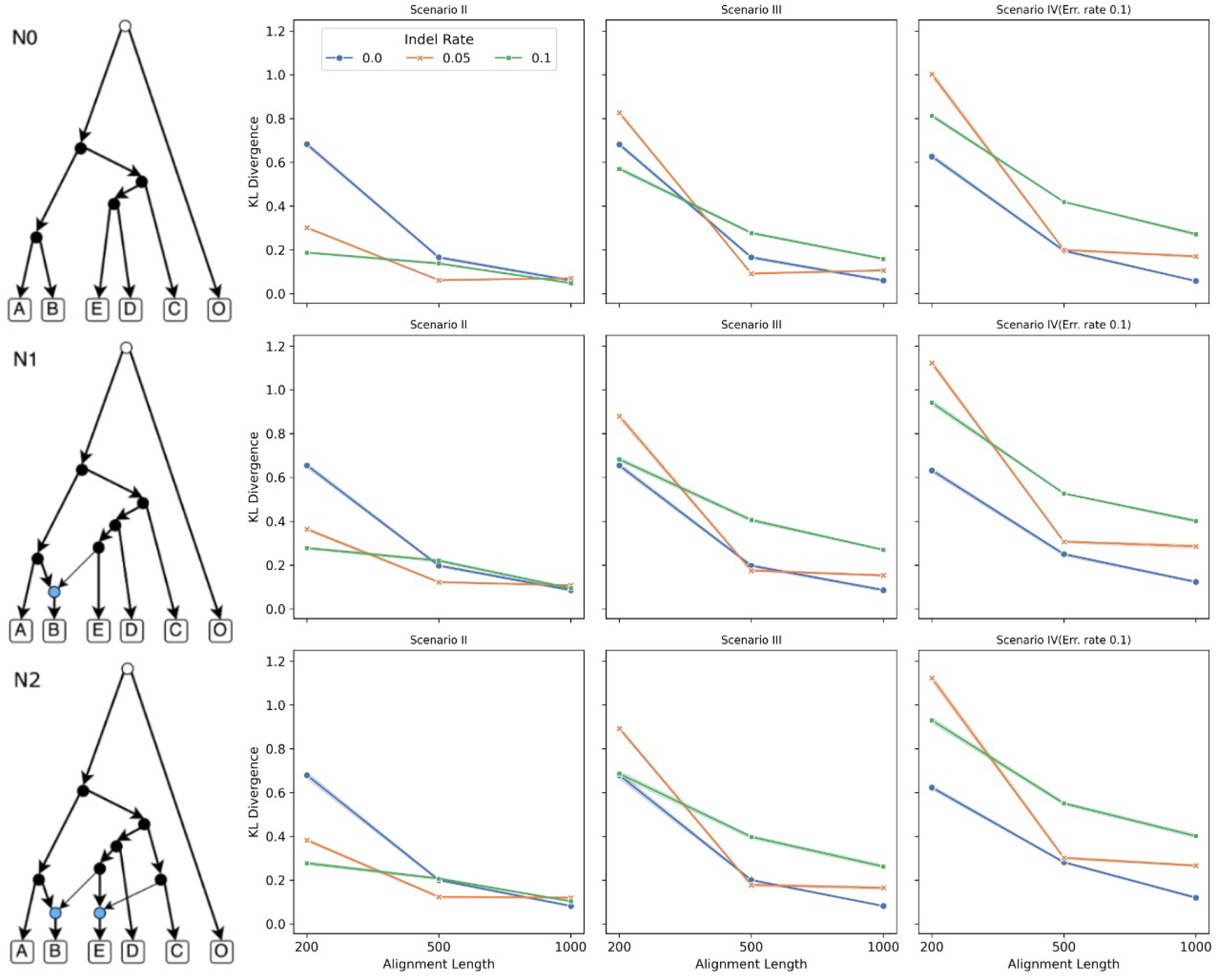
Changes in the distributions of gene tree topologies under the data error scenarios. The KL divergence between inferred and true gene tree topology distributions as a function of alignment length (x-axis), across different models (rows) and scenarios (columns). Each line represents a different indel rate.

For model N0 (top row), KL divergence consistently decreases with longer alignments across all scenarios and indel rates. In scenario II, the KL divergence at indel rate 0.0 appears unexpectedly high, but this is due to averaging across different numbers of gene trees: shorter alignments with only 200 gene trees show higher divergence, skewing the average upward. Scenario III follows a similar downward trend with alignment length but exhibits slightly higher divergence values than scenario II, especially at higher indel rates. Scenario IV yields the highest KL divergence overall, with divergence increasing clearly with indel rate—particularly for shorter alignments—highlighting the compounding effect of sequencing errors and indels.

Model N1 (middle row) shows patterns similar to N0, but with moderately higher KL divergence values overall, reflecting the greater complexity of the evolutionary history. Scenario IV again yields higher KL divergence than scenario III and II, particularly at high indel rates (0.1). Notably, scenario III shows a more substantial improvement with increasing alignment length than scenario IV, indicating that sequencing errors reduce the benefits of longer alignments.

Model N2 (bottom row) exhibits the largest KL divergence values across all scenarios, with scenario IV showing the steepest divergence at high indel rates. While KL values decrease with alignment length in all cases, the curves corresponding to high indel rates (orange and green) remain clearly separated from those with no indels (blue), especially in scenario IV. These results underscore the sensitivity of gene tree distribution fidelity to both indel rates and sequencing errors, particularly under more complex reticulation histories.

### 3.2 Likelihood trajectories with erroneous data

First, to assess how robust inference methods are to different sources of error, we evaluated absolute and relative changes in likelihood and pseudo-likelihood of the species networks inferred using the ML and MPL methods (Fig. 8–13).

**Figure 8:**
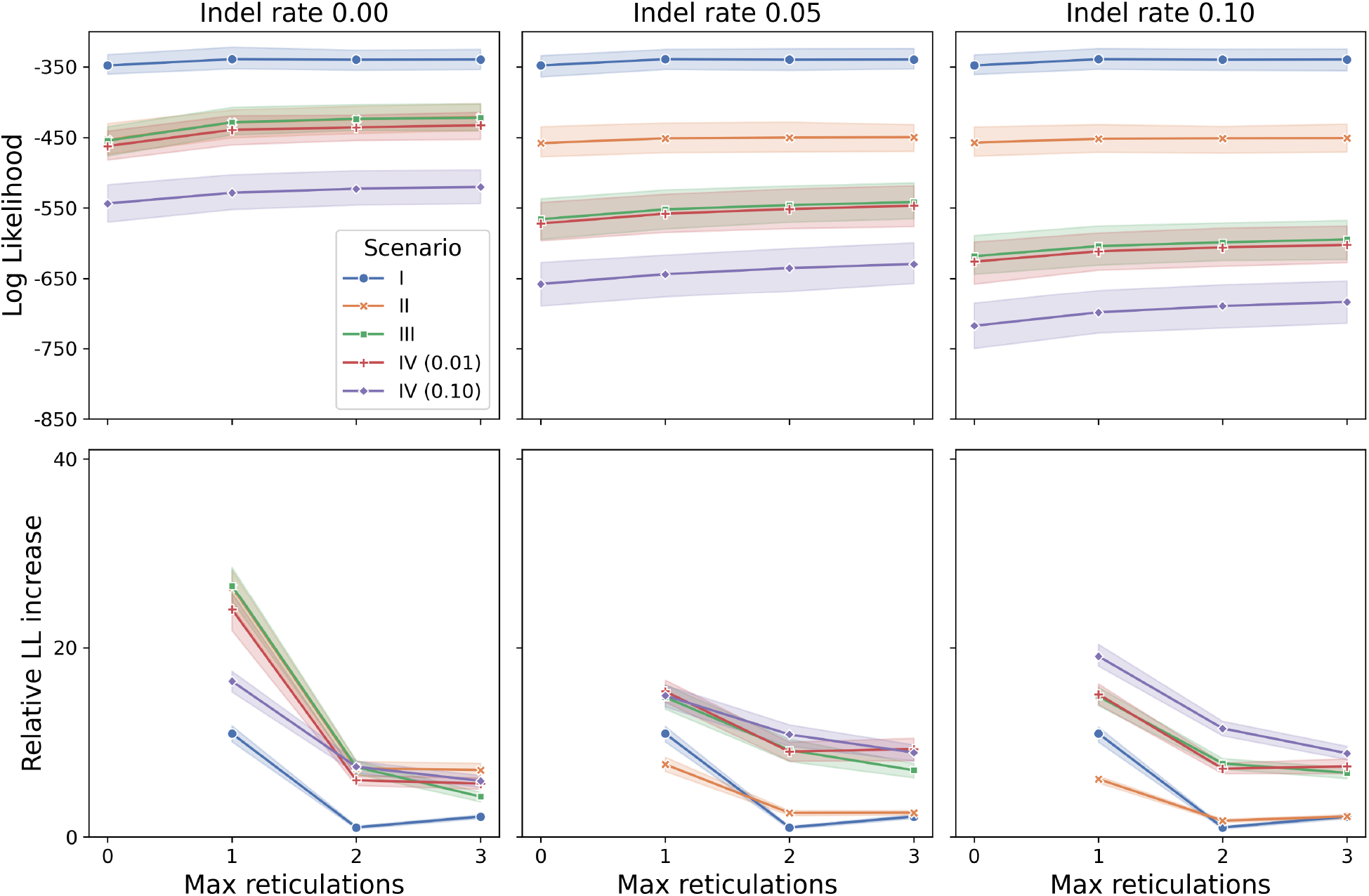
Likelihood and likelihood change for N0 species network in the presence of data error. **Top panels:** Log-likelihood of the MLE species network (ground truth: N0) as the function of the number of maximum reticulations allowed for candidate networks (x-axis). **Bottom panels:** Relative change in log-likelihood when the number of allowed reticulations increases by one, up to the maximum shown on the x-axis. In all subplots, colors correspond to inference scenarios, and in the case of scenario IV (modeling sequence error), the error rate setting.

#### Species tree inference (N0)

In this case, the ground truth is a species tree, so no reticulations need to be inferred. We observe that the MLE shows a slight upward trend in the likelihood of the inferred network as the number of allowed reticulations increases from 0 to 1 in scenario I (Fig. 8).

Additionally, in scenarios II, III, and IV—especially as the indel rate increases—allowing more reticulations continues to slightly improve the likelihood at each step (Fig. 8). This trend is most pronounced in scenario IV with 0.1 error rate, suggesting that overfitting to error increases with the level of error in the inferred gene trees.

Conversely, the MPLE shows no upward trend in scenario I (Fig. 9). Furthermore, in scenarios II, III, and IV, MPLE log pseudo-likelihood values remain remarkably stable.

**Figure 9:**
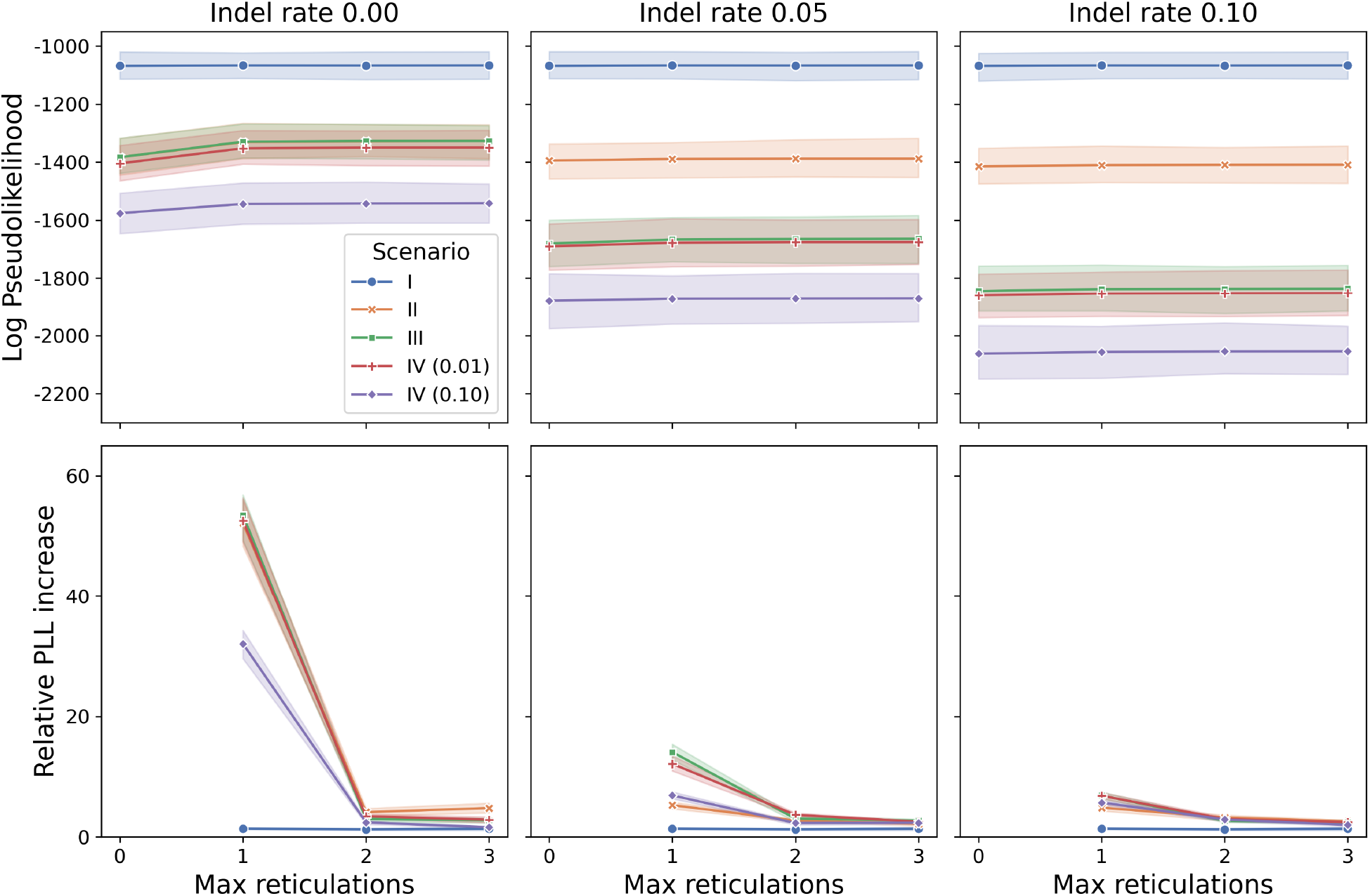
Pseudo-likelihood and pseudo-likelihood change for N0 species network in the presence of data error. **Top panels:** Log pseudo-likelihood of the MPLE species network (ground truth: N0) as the function of the number of maximum reticulations allowed for candidate networks (x-axis). **Bottom panels:** Relative change in log pseudo-likelihood when the number of allowed reticulations is increased by one, up to the maximum shown on the x-axis. In all subplots, colors correspond to inference scenarios, and in scenario IV, the error rate setting.

#### Species network inference with 1 reticulation (N1)

Here, the ground truth is a species network with one reticulation. The MLE exhibits a substantial increase in likelihood once the inference is allowed to include 1 reticulation (Fig. 10).

**Figure 10:**
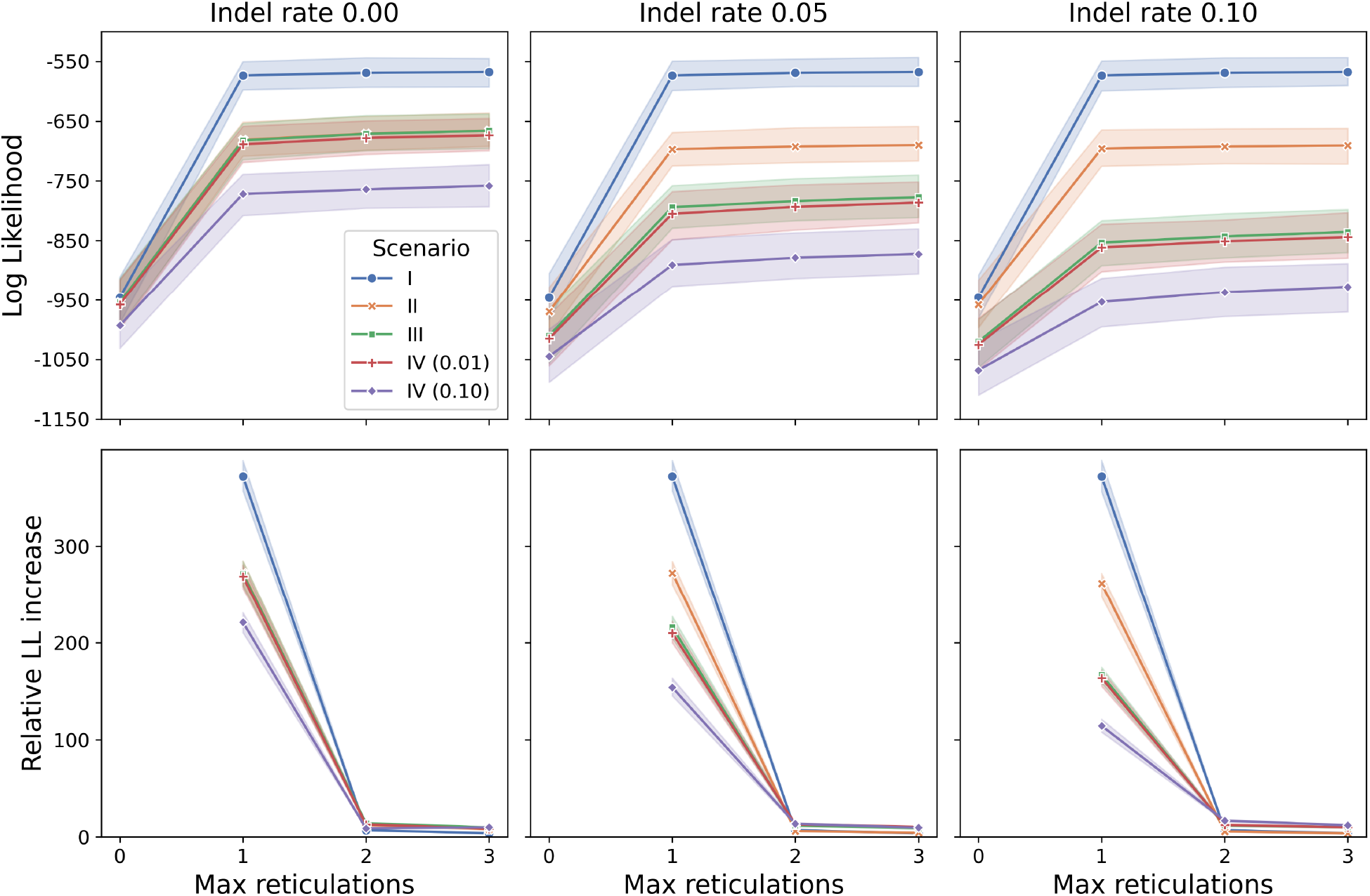
Likelihood and likelihood change for N1 species network in the presence of data error. **Top panels:** Log-likelihood of the MLE species network (ground truth: N1) as the function of the number of maximum reticulations allowed. **Bottom panels:** Relative change in log-likelihood when increasing the number of allowed reticulations by one. Colors correspond to inference scenarios, and in scenario IV, to the error rate.

Further increases in the number of allowed reticulations yield only minor improvements in likelihood (Fig. 10). As with the N0 case, the noisiest setting (scenario IV with 0.1 error rate) leads to the highest relative increase in likelihood from 1 to 2 reticulations allowed per network (Fig. 10, bottom panels).

Similarly to N0, MPLE is more stable than MLE between 1 and 3 allowed maximum reticulations (Fig. 11). There is no notable upward trend in the pseudo-likelihood even in scenario IV with 0.1 error rate. In fact, the relative pseudo-likelihood gain beyond 1 reticulation remains negligible in cases with non-zero indel rate (Fig. 11, bottom panels).

**Figure 11:**
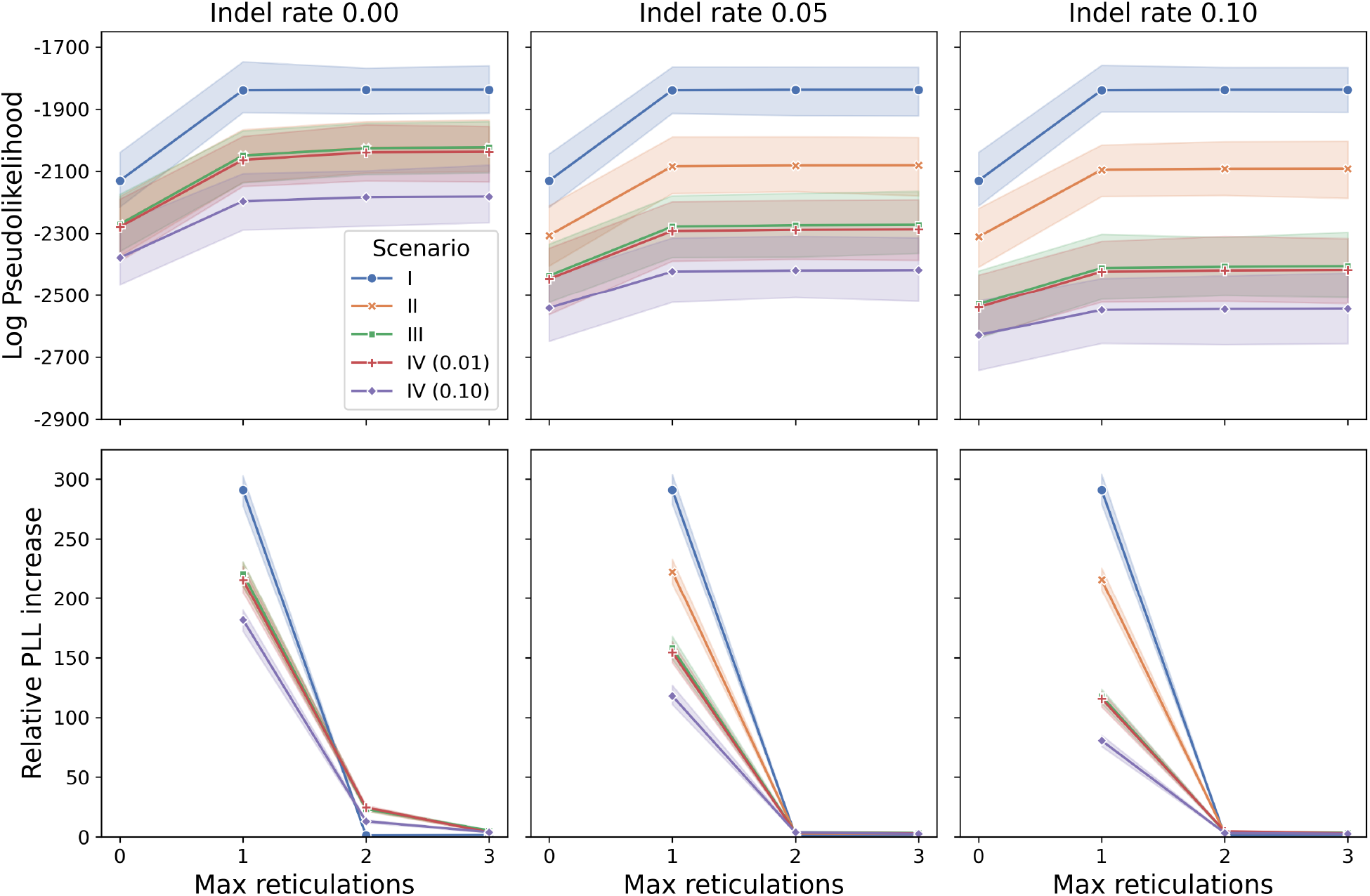
Pseudo-likelihood and pseudo-likelihood change for N1 species network in the presence of data error. **Top panels:** Log pseudo-likelihood of the MPLE species network (ground truth: N1) versus maximum reticulations allowed (x-axis). **Bottom panels:** Relative change in log pseudo-likelihood when increasing the number of allowed reticulations by one. In all subplots, the colors correspond to inference scenarios; for scenario IV, the colors also indicate the associated error rate.

#### Species network inference with 2 reticulations (N2)

Here, the ground truth includes two reticulations. As with N1, we see a large likelihood increase when more reticulations are allowed. However, the jump in likelihood from 1 to 2 reticulations being allowed is more modest (Fig. 12).

**Figure 12:**
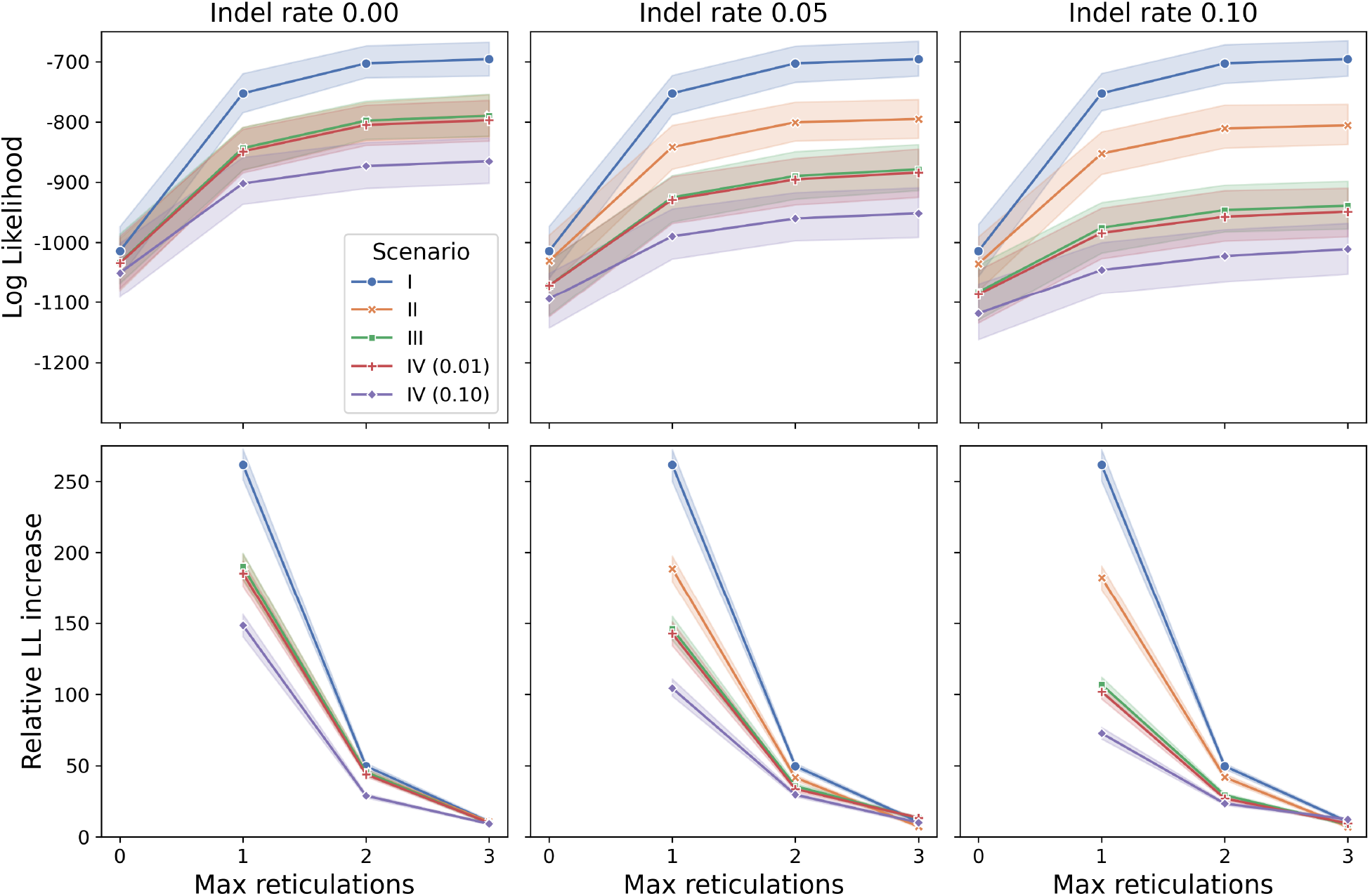
Likelihood and likelihood change for N2 species network in the presence of data error. **Top panels:** Log-likelihood of the MLE species network (ground truth: N2) versus maximum reticulations allowed (x-axis). **Bottom panels:** Relative change in log-likelihood when increasing the number of allowed reticulations by one. In all subplots, the colors correspond to inference scenarios; for scenario IV, the colors also indicate the associated error rate.

The relative log-likelihood gain as we allow up to 2 reticulations ranges from 25–50, which exceeds the relative increase at a similar point for the N1 case (Fig. 12, Fig. 10 bottom panels). Notably, in scenario IV with both error rate and indel rate at 0.1, the relative increase in likelihood resulting from allowing up to 3 reticulations is comparable to the 1-to-2 increase (Fig. 12 bottom right panel).

Analogously to the behavior observed for the N0 and N1 case, the MPLE shows greater stability than MLE from 2 to 3 allowed reticulations (Fig. 13). The relative pseudo-likelihood gain is nearly zero when increasing from 2 to 3 allowed reticulations.

**Figure 13:**
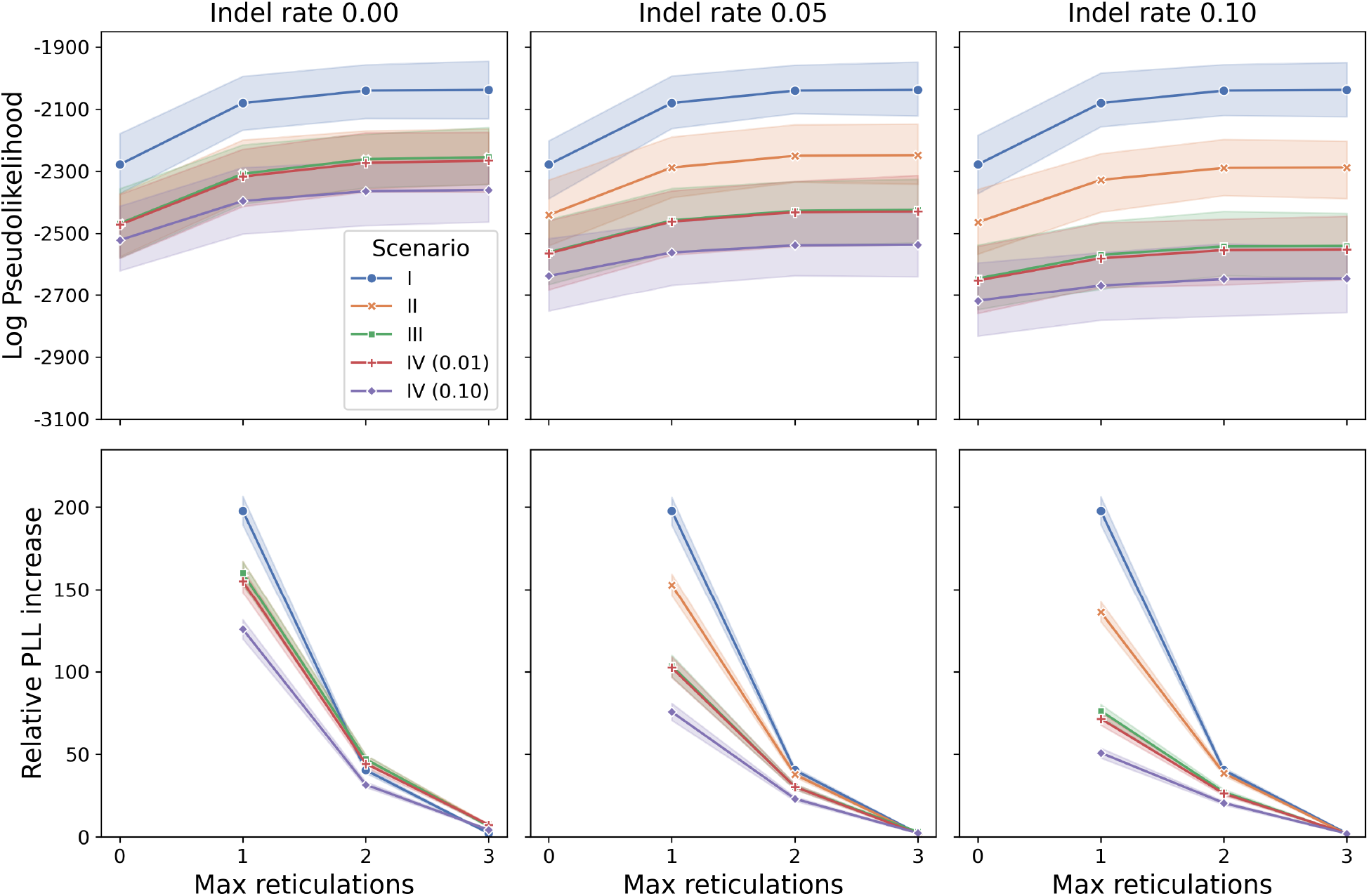
Pseudo-likelihood and pseudo-likelihood change for N2 species network in the presence of data error. **Top panels:** Log pseudo-likelihood of the MPLE species network (ground truth: N2) as the function of maximum reticulations (x-axis). **Bottom panels:** Relative change in log pseudo-likelihood as allowed reticulations increase by one. Scenario and error rate encodings are consistent with earlier figures.

### 3.3 Distributions of MCMC samples

In parallel with the likelihood and pseudo-likelihood trends observed in ML/MPL inference, we tracked the log pseudo-likelihood of the MAP estimates obtained from the MCMC runs (Fig. 14–16).

**Figure 14:**
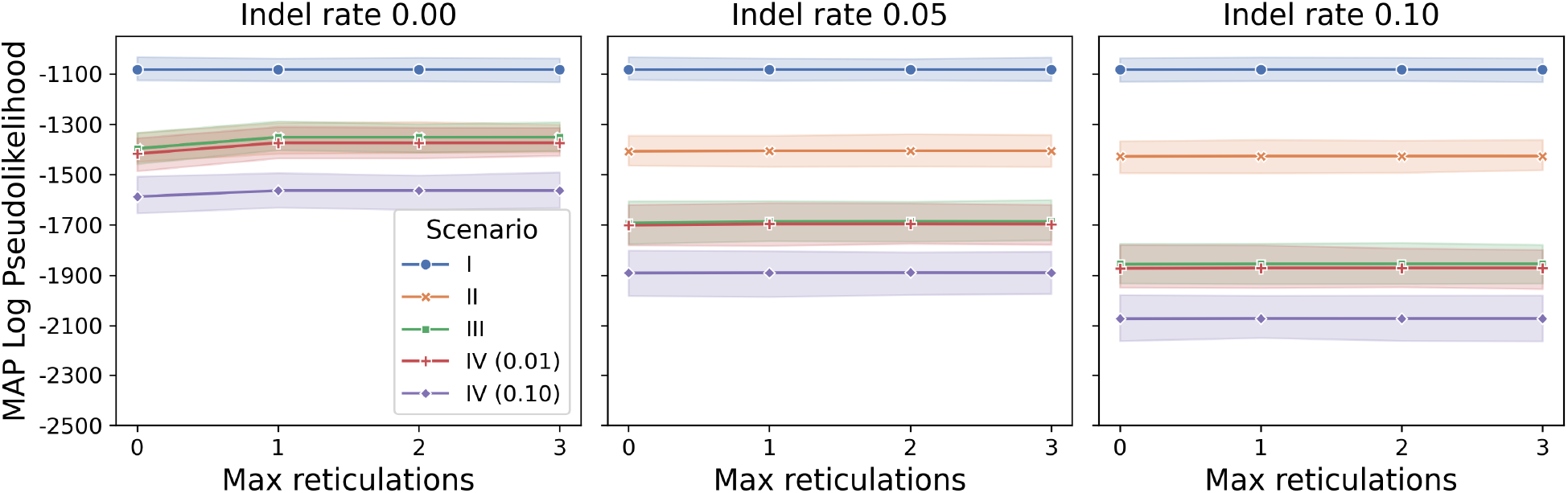
Pseudo-likelihood of the MAP estimate for the N0 species network in the presence of data error. Log pseudo-likelihood of the MAP species network estimate obtained from the MCMC run (ground truth: N0) as the function of the maximum allowed number of reticulations. The line shows the average value and the bands indicate the inter-quartile range.

For the N0 case, the pseudo-likelihood of the MAP estimates remained consistent as the number of allowed reticulations increased, with the only exception being a slight increase for data in scenarios II–IV without indels (Fig. 14).

In the N1 case, similar to the behavior observed with MPL inference, the pseudo-likelihood of the MAP estimate increased when one reticulation was allowed and remained relatively constant as the number of allowed reticulations increased further (Fig. 15).

**Figure 15:**
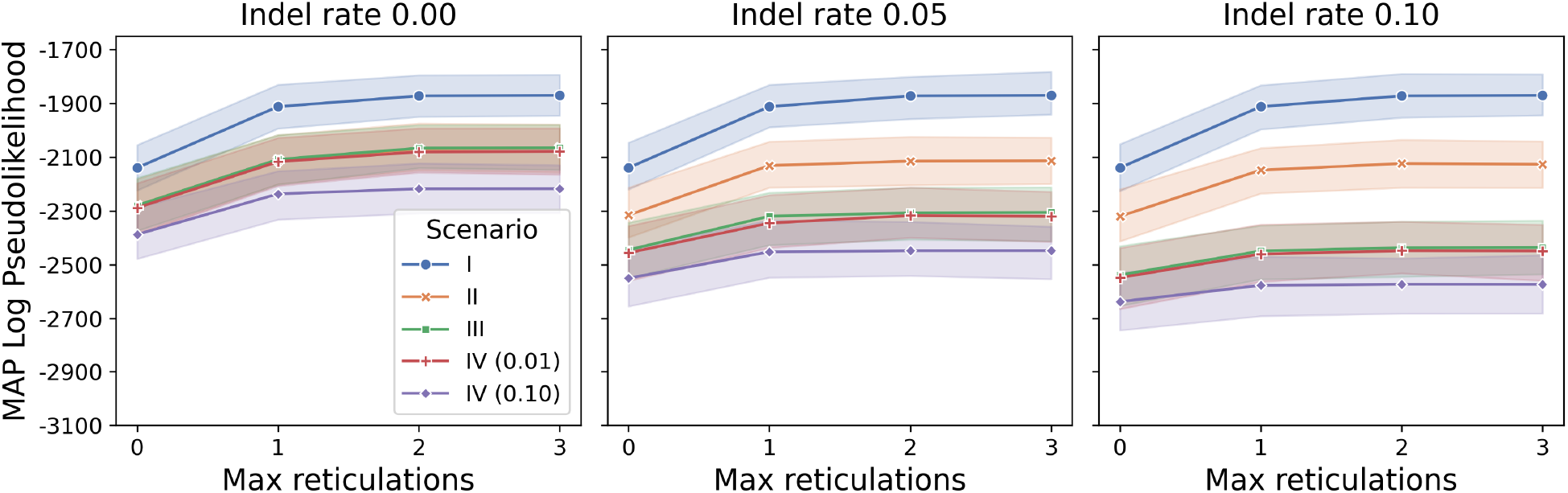
Pseudo-likelihood of the MAP estimate for the N1 species network in the presence of data error. Log pseudo-likelihood of the MAP species network estimate obtained from the MCMC run (ground truth: N1) as the function of the maximum allowed number of reticulations. The line shows the average value and the bands indicate the inter-quartile range.

For the N2 case, the pseudo-likelihood of the MAP estimates did not show a significant increase as the number of allowed reticulations went up from 1 to 2. This contrasts with the behavior observed in the ML and MPL inference results (Fig. 16, 13).

**Figure 16:**
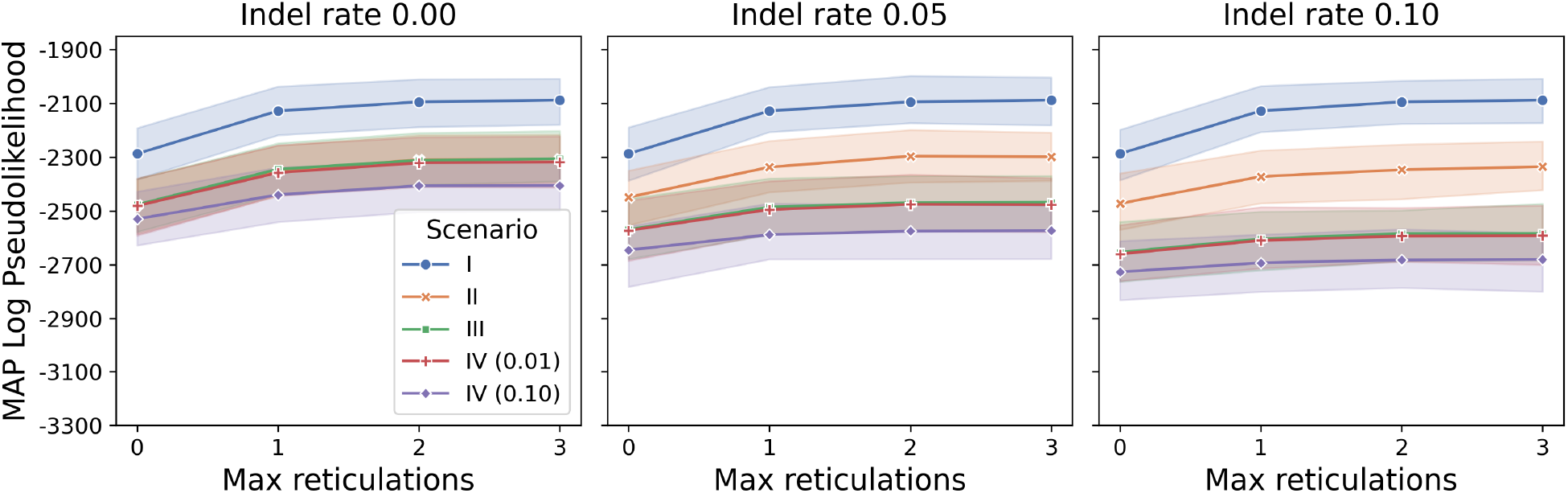
Pseudo-likelihood of the MAP estimate for the N2 species network in the presence of data error. Log pseudo-likelihood of the MAP species network estimate obtained from the MCMC run (ground truth: N2) as the function of the maximum allowed number of reticulations. The line shows the average value and the bands indicate the inter-quartile range.

### 3.4 Topological accuracy of inferred networks

We also evaluated topological error (Fig. 17) and topological accuracy (Fig. 18) to evaluate method performance on erroneous data. Due to the high number of parameter combinations, we focused on inference scenarios (I–IV; Fig. 3) and inference method (ML, MPL, MCMC), while visualizing variation due to indel rate, alignment length, and number of gene trees as inter-quartile ranges. We used the Robinson-Foulds distance [47] when both ground-truth and inferred topologies were trees, and the metric from [48] when at least one of the topologies was a network.

**Figure 17:**
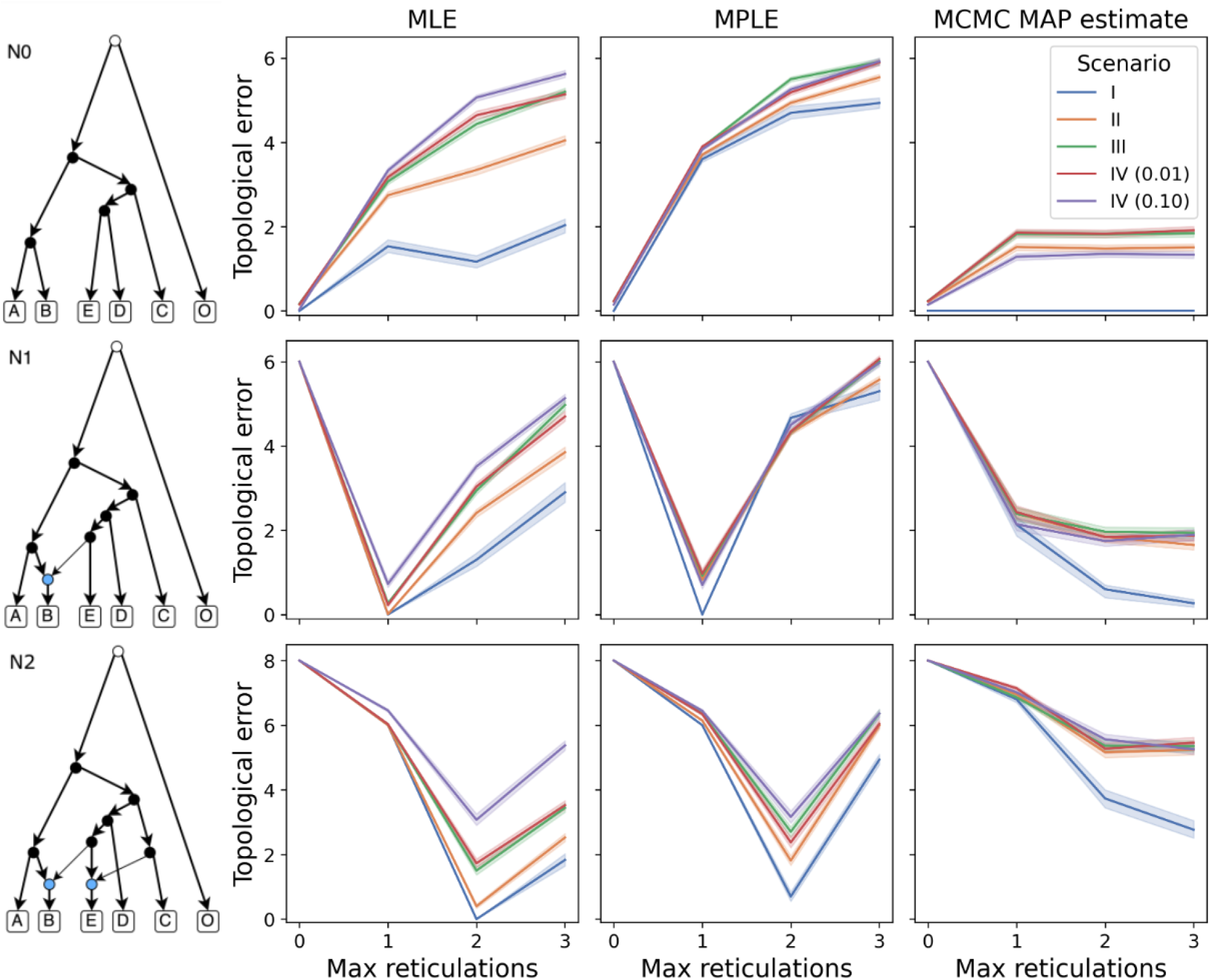
Topological error across inference methods and ground-truth species networks in the presence of data error. Each row shows the mean topological error for each method (ML, MPL, MCMC) across varied indel rates, alignment lengths, number of input gene trees, and replicate runs. Colored bands indicate inter-quartile ranges. The first row corresponds to N0 (species tree), the second to N1 (network with 1 reticulation), and the third to N2 (network with 2 reticulations). The x-axis denotes the maximum allowed number of reticulations, and colors indicate inference scenarios I–IV, with scenario IV further subdivided by error rates (0.01 and 0.1).

**Figure 18:**
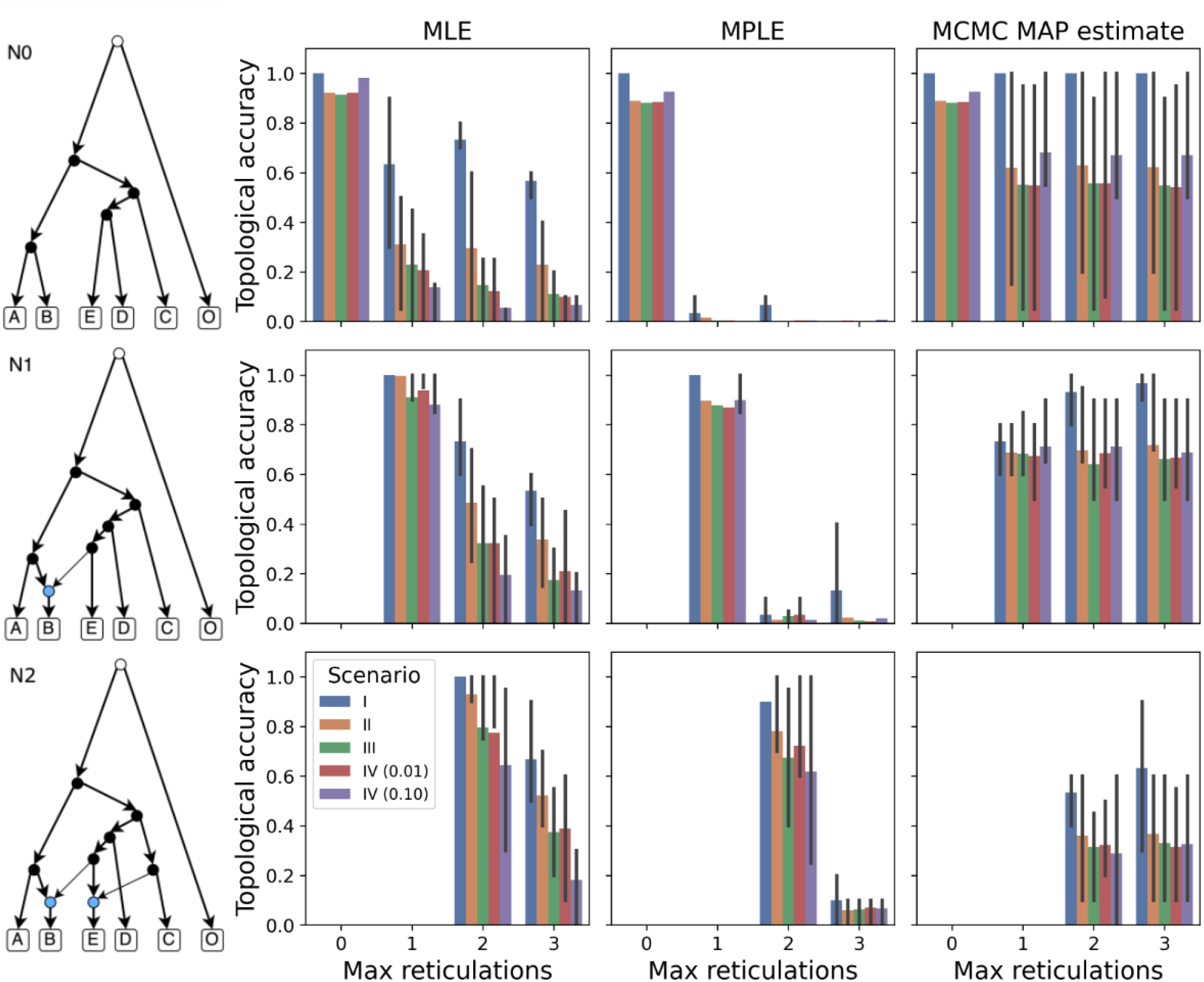
Topological accuracy across inference methods and ground-truth species networks in the presence of data error. Each row shows the mean number of correctly inferred topologies among 10 replicate runs for ML, MPL, and MCMC methods. Error bars represent inter-quartile ranges, reflecting variance due to indel rates, alignment lengths, and number of input gene trees. Rows correspond to N0, N1, and N2 networks, respectively. The x-axis indicates the maximum number of reticulations allowed, and colors indicate inference scenarios I–IV, with scenario IV further subdivided by error rates (0.01 and 0.1). Results are omitted when the number of allowed reticulations is less than the true value.

We found that both ML and MPL methods achieved the lowest error when the maximum number of allowed reticulations matched the true number (Fig. 17). In contrast, MCMC MAP estimates showed similar error across 1–3 maximum reticulations zone. Notably, all methods struggled in accurately recovering the topology of the N2 network under data error (Fig. 17).

Topological accuracy results show that ML and MPL methods perform well for N0 and N1 when the maximum number of reticulations equals the ground truth value, even in the presence of data error (Fig. 18). On the other hand, in the N1 and N2, MCMC MAP estimates were less accurate than ML and MPL when the number of reticulations allowed matches the true value but maintained comparable accuracy when allowed reticulations exceeded the ground-truth value (Fig. 18). Additionally, the MLE is the most affected by erroneous data, with accuracy decreasing as the complexity of the inference scenario increases (from I to IV (0.1)). Conversely, pseudo-likelihood-based methods (MPL and MCMC) are less sensitive to specific levels of error (Fig. 18), showing a notable change in accuracy primarily between scenario I (ground-truth gene trees) and the remaining scenarios (where gene trees are inferred).

### 3.5 Computational costs of inference

Error in the data has no impact on the computational cost of inference for methods based on heuristic search of the species network space. However, the three methods we evaluated differ substantially in runtime under their default settings, as shown in Table 2.

**Table 2:** CPU time required for running each of the evaluated inference methods on a single dataset. Columns represent mean runtime, standard deviation, and the 25th, 50th, and 75th percentiles. All times are in seconds.

| Method | CPU time (s) |  |  |  |  |
| --- | --- | --- | --- | --- | --- |
|  | Mean | St. dev. | 25% | 50% | 75% |
| InferNetwork_ML | 636.54 | 3169.78 | 77.15 | 145.69 | 371.96 |
| InferNetwork_MPL | 100.36 | 81.05 | 44.92 | 73.57 | 128.67 |
| MCMC_GT_Pseudo | 626.60 | 246.54 | 424.53 | 623.20 | 777.40 |

Among the methods, maximum-pseudo-likelihood inference is consistently the least resource-intensive method, with an average runtime of under two minutes. In contrast, Bayesian MCMC inference is the most computationally intensive, requiring nearly an order of magnitude more CPU time than MPL inference.

## 4 Discussion

In this study, we investigated the impact of data error on species network inference from gene tree data under the MSNC. Specifically, we examined how indels and sequencing errors affect alignment and gene tree inference, and how these propagate to distort gene tree distributions.

We found that indels have a more substantial effect on alignment accuracy and, consequently, on gene tree estimation error (GTEE) and shifts in gene tree distributions. This effect is particularly pronounced for shorter sequences, where indels can increase the average normalized RF distance between ground-truth and inferred gene trees by up to 0.1. While longer sequences mitigated this effect slightly, the maximum indel length was not scaled with sequence length, and hence the impact of indels might be underrepresented for longer sequences which can harbor larger indel events.

We also note that both maximum likelihood and maximum pseudo-likelihood inference methods perform well, and are partially robust to errors when the maximum number of reticulations allowed for the method matches the true number of reticulations in the species network. Allowing more reticulations often led to drop in accuracy, especially in the presence of error, as the models overfit to noise by introducing additional reticulations during the heuristic search. Notably, Bayesian inference under pseudo-likelihood via MCMC shows stronger performance in the presence of error even when the number of allowed reticulations exceeds the true number of reticulations. This is likely due to the default value of the prior for number of reticulations which is a Poisson distribution with parameter 1.

Although none of the evaluated methods can automatically identify the correct number of reticulations, tracking the (pseudo-)likelihood increase as a function of allowed reticulations provided useful signal about reasonable cutoff point. While error leads to overfitting and continued likelihood improvement with additional reticulations, the relative increase consistently goes down. Pseudo-likelihood, in particular, showed steeper declines in gain compared to full likelihood, especially when the ground truth was a species tree, suggesting it is less prone to overfitting due to its simpler modeling assumptions.

Taken together, these findings suggest that current network inference methods are reasonably robust to moderate data error and offer sufficient information for appropriate model choice. Maximum pseudolikelihood inference also offers the best computational efficiency, while showcasing some of the desired properties for distinguishing true reticulation events from overfitting to error, making it especially well-suited for large-scale genomic studies.

The most closely related prior study is that of Meijun *et al*. [31]. While their work focused on networks with a single reticulation and did not explore how data error interacts with inference complexity, our study systematically analyzed networks with up to two reticulations under a variety of error conditions—including alignment, sequencing, and gene tree estimation error. Unlike their approach, which didn’t infer the number of reticulations, we evaluated the behavior of likelihood and pseudo-likelihood values as a function of increasing reticulation complexity. This allowed us to demonstrate how overfitting arises in the presence of error and to identify inflection points in model fit. We also incorporated KL divergence to quantify how error distorts gene tree topology distributions—an aspect not covered in their analysis. Finally, our use of `MCMC GT Pseudo`, a pseudo-likelihood-based method that yields posterior distributions over networks and benefits from prior regularization. Together, these contributions provide a more comprehensive perspective on species network inference under data error and offer practical guidelines for interpreting network inference in empirical settings.

These results offer actionable recommendations for biologists applying species network inference to empirical datasets. First, the pronounced impact of indels and sequencing errors—especially in shorter alignments—highlights the importance of high-quality sequence assembly and alignment in minimizing downstream inference errors. Longer alignments or filtering by alignment quality and gene tree support can reduce noise. Second, since current tools do not estimate the number of reticulations automatically, practitioners are advised to evaluate changes in the (pseudo-)likelihood trajectory, using diminishing gain as a heuristic to avoid overfitting. Our findings also reinforce the value of pseudo-likelihood-based methods—such as `InferNetwork MPL` and `MCMC GT Pseudo`—for a good trade-off between computational efficiency and robustness to error, making them suitable for large-scale genomic analyses where data quality may vary.

Throughout our experiments, we used the single maximum likelihood estimate (MLE) gene tree per locus. To assess the reliability of these inferred gene trees and their agreement with the ground truth, bootstrap analysis is essential. For each replicate in the simulated dataset, we applied IQ-TREE’s ultrafast bootstrap method [49] with 1000 replicates to compute branch support values. Branches with bootstrap support below 0.7 were collapsed. From each collapsed gene tree, we extracted the set of bipartitions and compared them to those of the corresponding true gene tree. A collapsed gene tree was considered a refinement of the true gene tree if all of its bipartitions were present in the true tree. We then calculated the percentage of gene trees satisfying this refinement condition across replicates, reporting it as the *percentage of refinement*. The results are shown in Fig. 19.

**Figure 19:**
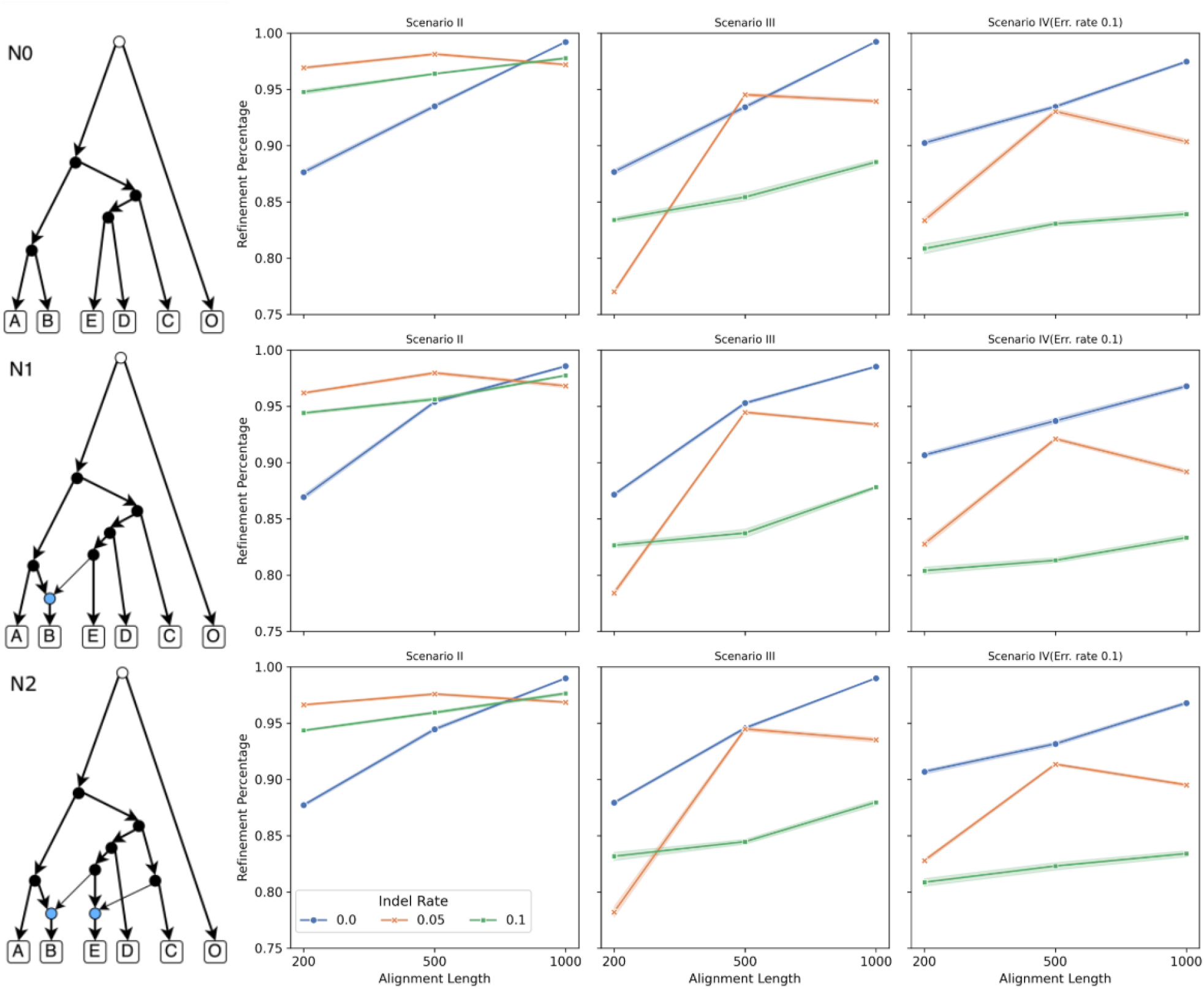
Bootstrap refinement of inferred gene trees under data error. Bootstrap-based refinement accuracy across different scenarios (columns) and evolutionary models (rows). Refinement is defined as the proportion of collapsed inferred gene trees whose bipartitions are a subset of the corresponding true gene tree. Collapsing was performed using a 0.7 bootstrap support threshold.

Each column allows a direct comparison of the effects of alignment error and sequencing error on refinement accuracy. In scenario II, we observe the highest refinement percentages across all models, indicating that gene tree inference is most accurate when alignments are error-free. In scenario III, refinement percentages decline slightly, reflecting the impact of alignment ambiguity. Scenario IV exhibits the lowest refinement percentages in all models. A closer comparison between scenario III and scenario IV further highlights the effect of sequencing error in the presence of varying indel rates. For moderate (0.05, orange) and high (0.1, green) indel rates, refinement percentages drop more substantially in scenario IV than in scenario III, particularly for shorter alignments. This pattern implies that sequencing error and indels interact synergistically to reduce inference accuracy, exacerbating the disruption caused by each individually. These effects are consistent across all model conditions (N0–N2).

These results indicate that one potential solution for dealing with gene tree error in practice is to first collapse poorly supported branches in the gene tree estimates, and then allow the species tree/network inference method to operate on the resolutions of the resulting non-binary gene trees. Indeed, this option is implemented in several likelihood and parsimony inference methods in `PhyloNet` and has been shown to be a promising approach in the context of speeding up Bayesian MCMC inference from sequence alignment data [50]. Furthermore, while approaches have been proposed to “fix” gene trees with respect to species trees, a recent study has highlighted the downsides of such strategies [51]. In general, enabling statistical or summary methods to work with non-binary gene trees and incorporate their refinements significantly increase computational cost compared to using single point estimates.

There are several limitations to our current study that should be addressed in future work. First, while we explored the impact of sequence errors on alignment and gene tree estimation, as well as the overall influence of data error on species network inference, we focused only on single-nucleotide errors. The impact of more complex forms of error—such as indels and short repeat expansions or contractions—remains to be evaluated. Moreover, beyond sequence-level errors, inaccuracies due to incorrect haplotyping introduce a different set of challenges. Although some preliminary studies have investigated the effects of haplotype phasing on reticulation detection [11, 52], more comprehensive analyses are necessary.

Second, our evaluation centered on inference methods that operate from estimated gene trees. As a result, all sources of data error ultimately propagate into distortions in gene tree topology distributions. Other methods in `PhyloNet`, such as `MCMC SEQ` [12], which infers species networks directly from sequence alignments, and `MLE BiMarkers` [15] and `MCMC BiMarkers` [13], which infer from bi-allelic markers, work directly with sequence data, rather than gene tree estimates. We expect that these methods—especially those using bi-allelic markers—may exhibit increased sensitivity to data error, and further investigation is required to fully understand the implications for inference.

Third, our simulations followed the MSNC model and assumed a single sampled individual per species. A more extensive evaluation should examine how gene duplication and loss events, as well as sampling multiple individuals within species, influence the robustness and accuracy of network inference.

## Acknowledgments

This work was in part supported by the NSF grants DMS/NIGMS-2153704 and DBI-2030604.

## Footnotes

1 The term ‘network’ is used to mean many things in biology. In this chapter, we use ‘species network’ (and, simply just ‘network’), as the counterpart to ‘species tree,’ referring specifically to the explicit phylogenetic network of a set of species.

## References

[1] Sebastien Roch and Sagi Snir. Recovering the treelike trend of evolution despite extensive lateral genetic transfer: A probabilistic analysis. Journal of Computational Biology, 20:93–112, 2013. doi: 10.1089/cmb.2012.0234.

[2] Ruth Davidson, Pranjal Vachaspati, Siavash Mirarab, and Tandy Warnow. Phylogenomic species tree estimation in the presence of incomplete lineage sorting and horizontal gene transfer. BMC Genomics, 16:S1, 2015. doi: 10.1186/1471-2164-16-S10-S1.

[3] Claudia Solís-Lemus and Cécile Ané. Inferring phylogenetic networks with maximum pseudolikelihood under incomplete lineage sorting. PLoS Genetics, 12:e1005896, 2016. doi: 10.1371/journal.pgen.1005896.

[4] Jiafan Zhu, Yun Yu, and Luay Nakhleh. In the light of deep coalescence: Revisiting trees within networks. BMC Bioinformatics, 17:415, 2016. doi: 10.1186/s12859-016-1269-1.

[5] Zhen Cao, Xinhao Liu, Huw A Ogilvie, Zhi Yan, and Luay Nakhleh. Practical aspects of phylogenetic network analysis using phyloNet. In Laura S Kubatko and L Lacey Knowles, editors, Species Tree Inference, pages 89–119. Princeton University Press, 2023. doi: 10.1515/9780691245157.

[6] Luay Nakhleh. Evolutionary Phylogenetic Networks: Models and Issues, pages 125–158. Springer US, Boston, MA, 2011. ISBN 978-0-387-09760-2. doi: 10.1007/978-0-387-09760-2\_7.

[7] James H Degnan and Noah A Rosenberg. Gene tree discordance, phylogenetic inference and the multispecies coalescent. Trends in Ecology & Evolution, 24:332–340, 2009. doi: 10.1016/j.tree.2009.01.009.

[8] Yun Yu, James H Degnan, and Luay Nakhleh. The probability of a gene tree topology within a phylogenetic network with applications to hybridization detection. PLoS Genetics, 8:e1002660, 2012.

[9] Yun Yu, Jianrong Dong, Kevin Liu, and Luay Nakhleh. Maximum likelihood inference of reticulate evolutionary histories. Proceedings of the National Academy of Sciences, 111(46):16448–16453, 2014. doi: 10.1073/pnas.1407950111.

[10] Yun Yu, R Matthew Barnett, and Luay Nakhleh. Parsimonious inference of hybridization in the presence of incomplete lineage sorting. Systematic Biology, 62:738–751, 2013. doi: 10.1093/sysbio/syt037.

[11] Dingqiao Wen, Yun Yu, and Luay Nakhleh. Bayesian inference of reticulate phylogenies under the multispecies network coalescent. PLOS Genetics, 12:e1006006, 2016. doi: 10.1371/journal.pgen.1006006.

[12] Dingqiao Wen and Luay Nakhleh. Co-estimating reticulate phylogenies and gene trees from multi-locus sequence data. Systematic Biology, 67:439–457, 2018. doi: 10.1093/sysbio/syx085.

[13] Jiafan Zhu, Dingqiao Wen, Yun Yu, Heidi M. Meudt, and Luay Nakhleh. Bayesian inference of phylogenetic networks from bi-allelic genetic markers. PLOS Computational Biology, 14:e1005932, 2018. doi: 10.1371/journal.pcbi.1005932.

[14] Chi Zhang, Huw A Ogilvie, Alexei J Drummond, and Tanja Stadler. Bayesian inference of species networks from multilocus sequence data. Molecular Biology and Evolution, 35:504–517, 2018. doi: 10.1093/molbev/msx307.

[15] Jiafan Zhu and Luay Nakhleh. Inference of species phylogenies from bi-allelic markers using pseudo-likelihood. Bioinformatics, 34:i376–i385, 2018. doi: 10.1093/bioinformatics/bty295.

[16] Tomáš Flouri, Xiyun Jiao, Bruce Rannala, and Ziheng Yang. A Bayesian implementation of the multispecies coalescent model with introgression for phylogenomic analysis. Molecular Biology and Evolution, 37:1211–1223, 2020. doi: 10.1093/molbev/msz296.

[17] RA Leo Elworth, Huw A Ogilvie, Jiafan Zhu, and Luay Nakhleh. Advances in computational methods for phylogenetic networks in the presence of hybridization. In Bioinformatics and Phylogenetics, pages 317–360. Springer, 2019.

[18] Siavash Mirarab, Luay Nakhleh, and Tandy Warnow. Multispecies coalescent: Theory and applications in phylogenetics. Annual Review of Ecology, Evolution, and Systematics, 52(Volume 52, 2021):247–268, 2021. ISSN 1545-2069. doi: 10.1146/annurev-ecolsys-012121-095340.

[19] Huw A Ogilvie, Remco R Bouckaert, and Alexei J Drummond. Starbeast2 brings faster species tree inference and accurate estimates of substitution rates. Molecular Biology and Evolution, 34(8): 2101–2114, 2017.

[20] David Bryant, Remco Bouckaert, Joseph Felsenstein, Noah A Rosenberg, and Arindam RoyChoudhury. Inferring species trees directly from biallelic genetic markers: bypassing gene trees in a full coalescent analysis. Molecular biology and evolution, 29(8):1917–1932, 2012.

[21] Charles-Elie Rabier, Vincent Berry, Jean-Christophe Glaszmann, Fabio Pardi, and Céline Scornavacca. On the inference of complex phylogenetic networks by markov chain monte-carlo. bioRxiv, 2020.

[22] Eric Y. Durand, Nick Patterson, David Reich, and Montgomery Slatkin. Testing for ancient admixture between closely related populations. Molecular Biology and Evolution, 28(8):2239–2252, 2011. doi: 10.1093/molbev/msr048.

[23] RA Leo Elworth, Chabrielle Allen, Travis Benedict, Peter Dulworth, and Luay Nakhleh. ???????? : A test statistic for detection of general introgression scenarios. In Proceedings of the 18th Workshop on Algorithms in Bioinformatics (WABI), 2018.

[24] James B Pease and Matthew W Hahn. Detection and polarization of introgression in a five-taxon phylogeny. Systematic biology, 64(4):651–662, 2015.

[25] Paul D Blischak, Julia Chifman, Andrea D Wolfe, and Laura S Kubatko. HyDe: A Python package for genome-scale hybridization detection. Systematic Biology, 67:821–829, 2018. doi: 10.1093/sysbio/syy023.

[26] Laura S. Kubatko and Julia Chifman. An invariants-based method for efficient identification of hybrid species from large-scale genomic data. BMC Evolutionary Biology, 19:112, 2019. doi: 10.1186/s12862-019-1439-7.

[27] Yun Yu and Luay Nakhleh. A maximum pseudo-likelihood approach for phylogenetic networks. BMC genomics, 16(10):1–10, 2015.

[28] Zhen Cao, Meng Li, Huw Ogilvie, and Luay Nakhleh. The impact of model misspecification on phylogenetic network inference. Bulletin of the Society of Systematic Biologists, 3(1), Sep. 2024. doi: 10.18061/bssb.v3i1.9553.

[29] Claudia Solís-Lemus, Mengyao Yang, and Cécile Ané. Inconsistency of species tree methods under gene flow. Systematic Biology, 65:843–851, 2016. doi: 10.1093/sysbio/syw030.

[30] Jun Huang, Yuttapong Thawornwattana, Tomáš Flouri, James Mallet, and Ziheng Yang. Inference of gene flow between species under misspecified models. Molecular Biology and Evolution, 39(12): msac237, 2022.

[31] Meijun Gao, Wei Wang, and Kevin J Liu. The impact of gene sequence alignment and gene tree estimation error on summary-based species network estimation. In Proceedings of the 13th ACM International Conference on Bioinformatics, Computational Biology and Health Informatics, pages 1–17, 2022.

[32] Daniel H Huson and David Bryant. Application of phylogenetic networks in evolutionary studies. Molecular biology and evolution, 23(2):254–267, 2006.

[33] Yun Yu, Cuong Than, James H. Degnan, and Luay Nakhleh. Coalescent histories on phylogenetic networks and detection of hybridization despite incomplete lineage sorting. Systematic Biology, 60(2): 138–149, 2011.

[34] Joseph Felsenstein. Evolutionary trees from DNA sequences: a maximum likelihood approach. Journal of Molecular Evolution, 17(6):368–376, 1981.

[35] Jiafan Zhu, Xinhao Liu, Huw A Ogilvie, and Luay K Nakhleh. A divide-and-conquer method for scalable phylogenetic network inference from multilocus data. Bioinformatics, 35(14):i370–i378, 2019.

[36] Nicolae Sapoval, Zejian Liu, Mehrdad Tamiji, Meng Li, and Luay Nakhleh. Theoretical and empirical performance of pseudo-likelihood-based bayesian inference of species trees under the multispecies coalescent. bioRxiv, pages 2025–01, 2025.

[37] Cuong Than, Derek Ruths, and Luay Nakhleh. Phylonet: A software package for analyzing and reconstructing reticulate evolutionary relationships. BMC Bioinformatics, 9:322, 2008. doi: 10.1186/1471-2105-9-322.

[38] Dingqiao Wen, Yun Yu, Jiafan Zhu, and Luay Nakhleh. Inferring phylogenetic networks using PhyloNet. Systematic Biology, 67:735–740, 2018. doi: 10.1093/sysbio/syy015.

[39] Cuong Than and Luay Nakhleh. Species tree inference by minimizing deep coalescences. PLoS computational biology, 5(9):e1000501, 2009.

[40] Richard R Hudson. Generating samples under a Wright-Fisher neutral model of genetic variation. Bioinformatics, 18:337–338, 2002. doi: 10.1093/bioinformatics/18.2.337.

[41] William Fletcher and Ziheng Yang. Indelible: a flexible simulator of biological sequence evolution. Molecular biology and evolution, 26(8):1879–1888, 2009.

[42] Kazutaka Katoh and Daron M Standley. Mafft multiple sequence alignment software version 7: improvements in performance and usability. Molecular biology and evolution, 30(4):772–780, 2013.

[43] Bui Quang Minh, Heiko A Schmidt, Olga Chernomor, Dominik Schrempf, Michael D Woodhams, Arndt Von Haeseler, and Robert Lanfear. IQ-TREE 2: new models and efficient methods for phylogenetic inference in the genomic era. Molecular biology and evolution, 37(5):1530–1534, 2020.

[44] Mehrdad Tamiji, Nicolae Sapoval, and Luay Nakhleh. Data from “Impact of Data Error on Phylogenetic Network Inference from Gene Trees Under the Multispecies Network Coalescent”, 2025. URL 10.5281/zenodo.15345227.

[45] Mehrdad Tamiji, Nicolae Sapoval, and Luay Nakhleh. Scripts for “Impact of Data Error on Phylogenetic Network Inference from Gene Trees Under the Multispecies Network Coalescent”, 2025. URL https://github.com/nakhlehlab/data-error-species-nets/.

[46] PhyloNet. URL https://github.com/NakhlehLab/PhyloNet.

[47] David F Robinson and Leslie R Foulds. Comparison of phylogenetic trees. Mathematical Biosciences, 53:131–147, 1981. doi: 10.1016/0025-5564(81)90043-2.

[48] Luay Nakhleh. A metric on the space of reduced phylogenetic networks. IEEE/ACM Transactions on Computational Biology and Bioinformatics, 7:218–222, 2009. doi: 10.1109/TCBB.2009.2.

[49] Diep Thi Hoang, Olga Chernomor, Arndt von Haeseler, Bui Quang Minh, and Le Sy Vinh. UFBoot2: improving the ultrafast bootstrap approximation. Molecular Biology and Evolution, 35(2):518–522, 10 2017. ISSN 0737-4038. doi: 10.1093/molbev/msx281.

[50] Yaxuan Wang, Huw A Ogilvie, and Luay Nakhleh. Practical speedup of bayesian inference of species phylogenies by restricting the space of gene trees. Molecular biology and evolution, 37(6):1809–1818, 2020.

[51] Zhi Yan, Huw A Ogilvie, and Luay Nakhleh. “correcting” gene trees to be more like species trees frequently increases topological error. Genome Biology and Evolution, 15(6):evad094, 2023.

[52] Jun Huang, Jeremy Bennett, Tomáš Flouri, Adam D Leaché, and Ziheng Yang. Phase resolution of heterozygous sites in diploid genomes is important to phylogenomic analysis under the multispecies coalescent model. Systematic Biology, 71(2):334–352, 2022.

